# Novel incursion of a highly pathogenic avian influenza subtype H5N8 virus in the Netherlands, October 2020

**DOI:** 10.1101/2020.11.03.361634

**Authors:** Nancy Beerens, Rene Heutink, Frank Harders, Marit Roose, Sylvia B.E. Pritz-Verschuren, Evelien A. Germeraad, Marc Engelsma

**Affiliations:** Wageningen Bioveterinary Research, Lelystad, the Netherlands

**Keywords:** HPAI H5N8, genome sequence, evolution, swans, wild birds

## Abstract

The HPAI H5N8 virus detected in mute swans in the Netherlands in October 2020 shares a common ancestor with clade 2.3.4.4b viruses last detected in Egypt in 2018-2019 and has a similar genetic composition. The virus is not directly related to European H5N8 viruses detected in first half of 2020.

---

The introduction of highly pathogenic avian influenza (HPAI) H5 clade 2.3.4.4 viruses in Europe had large impact on animal health and caused substantial losses to the poultry industry between 2014 and 2020. Migratory waterfowl are implicated in the distribution of HPAI H5 viruses along flyways from breeding grounds in North-Russia to wintering sites in Europe (1, 2). In 2016, HPAI H5N8 viruses of clade 2.3.4.4b were introduced in Europe, including the Netherlands, infecting several poultry farms and causing high mortality amongst wild birds (3). Analysis of wild bird and poultry viruses revealed several independent incursion events and frequent reassortment events (4). In the first months of 2020, HPAI H5N8 clade 2.3.4.4b viruses were detected in Eastern-Europe and Germany (5, 6). These HPAI H5N8 viruses were genetically distinct from the viruses detected in Europe in 2016. Early in 2020 also HPAI H5N8 viruses were detected in Bulgaria (6). In October 2020 HPAI H5N8 virus was detected in wild birds in the Netherlands. In this study, we performed genetic and phylogenetic analysis to study the origin of this virus, and its relationship with viruses previously detected in Europe.

## The Study

Two mute swans (*Cygnus olor*) were found dead in the Province of Utrecht, located in the central part of the Netherlands. The mute swans were submitted for diagnostic testing as part of the wild bird surveillance program for avian influenza virus. Swab samples from trachea and cloaca tested positive in the screening M-PCR *(7)* and H5-PCR *(8)*, demonstrating a notifiable subtype H5 virus. Hemagglutinin (HA) and neuraminidase (NA) sequence analysis was performed as previously described *(7)*. The HA cleavage site PLREKRRKR*GLF showed polybasic properties characteristic of highly pathogenic viruses, and NA sequence results subtyped the virus as H5N8. Full genome sequencing was performed using the Illumina MiSeq-platform, as described previously (4). Database searches showed that the virus (A/mute_swan/Netherlands/20015931-001/2020; EPI591075) is related to HPAI H5N8 clade 2.3.4.4b viruses previously detected in Eurasia in 2016-2018 for all gene segments (Table 1). For six of the gene segments, a virus isolated in Greonterp (the Netherlands) in 2016 was identified as highly similar, with a nucleotide homology between 97%-98%. For the matrix (MP) segment, viruses with higher nucleotide homology (99%) were detected in Asia and Egypt in 2016-2017.

**Table 1:**
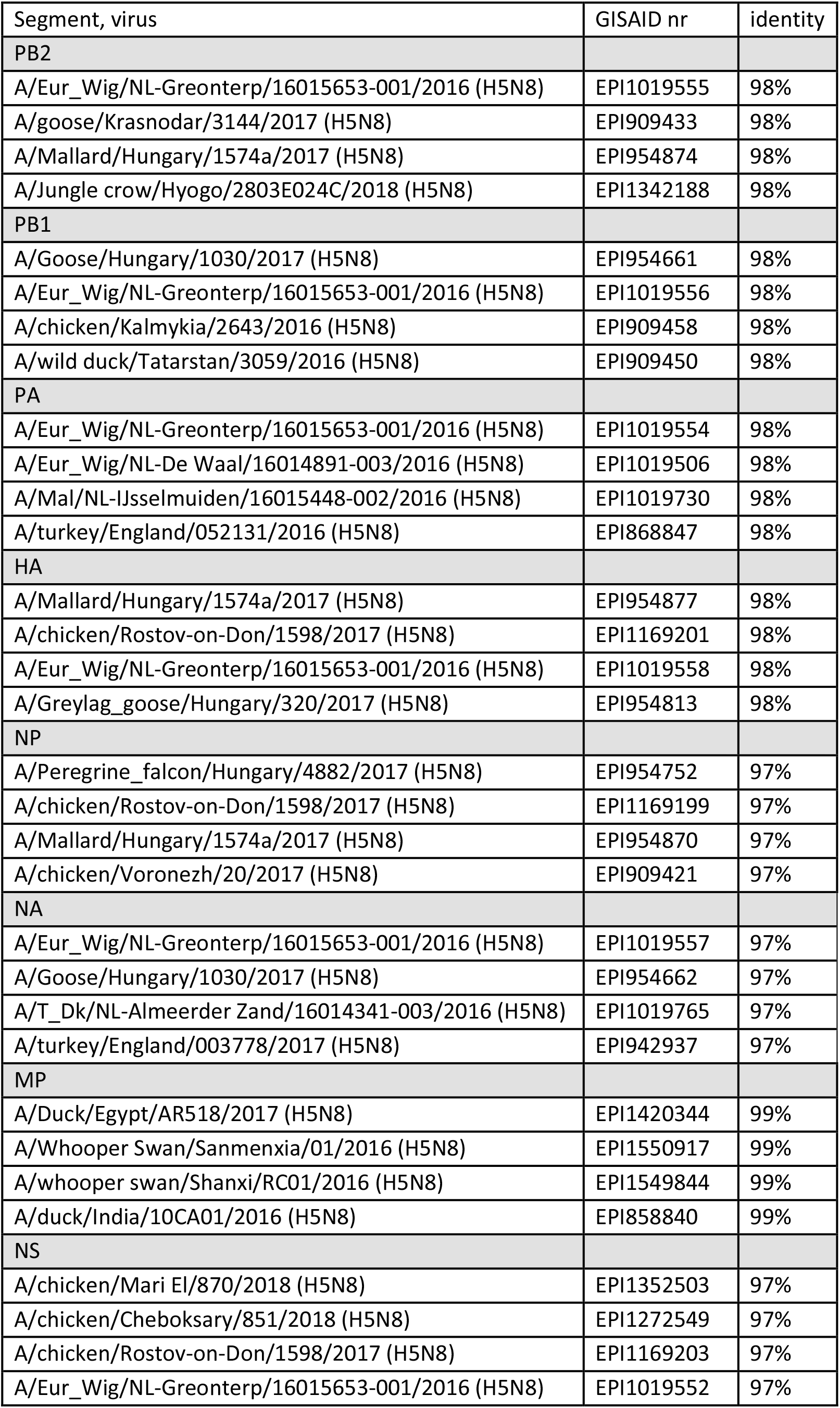
Genetic composition of the HPAI H5N8 virus isolated in the Netherlands, 2020

The genetic relationship between the HPAI H5N8 viruses detected in the Netherlands in 2016 and 2020 raised questions on possible long-term undetected circulation of the virus in local wild bird populations. To study the origin of the novel H5N8 virus, we performed detailed phylogenetic analysis for all gene segments. Phylogenetic analysis of the HA (Fig 1) and NA (Supplementary FigS1) shows that the closest genetic relative was isolated from a duck in Egypt in January 2019 (EPI399644, only HA and NA sequences publicly available). The virus also shares a common ancestor with other viruses detected in Egypt in 2018-2019, and with viruses detected in the Netherlands and Eurasia in 2016-2017. The novel H5N8 virus does not cluster with the H5N8 viruses that caused widespread outbreaks in Eastern-Europe and Germany earlier in 2020, or viruses detected in Bulgaria in 2020. For all other gene segments of the novel H5N8 virus, clustering was observed with H5N8 viruses that circulated in Egypt in 2018-2019 and in Eurasia in 2016-2017 (Supplementary FigS1). However, several viruses closely related to MP were found to be more distantly related for the other gene segments which suggest reassortment has occurred for the MP segment. HPAI H5N8 viruses closely related to MP were isolated in Asia and Egypt in 2016-2018. No reassortments with low pathogenic avian influenza viruses were observed for any of the segments.

**Fig 1.**
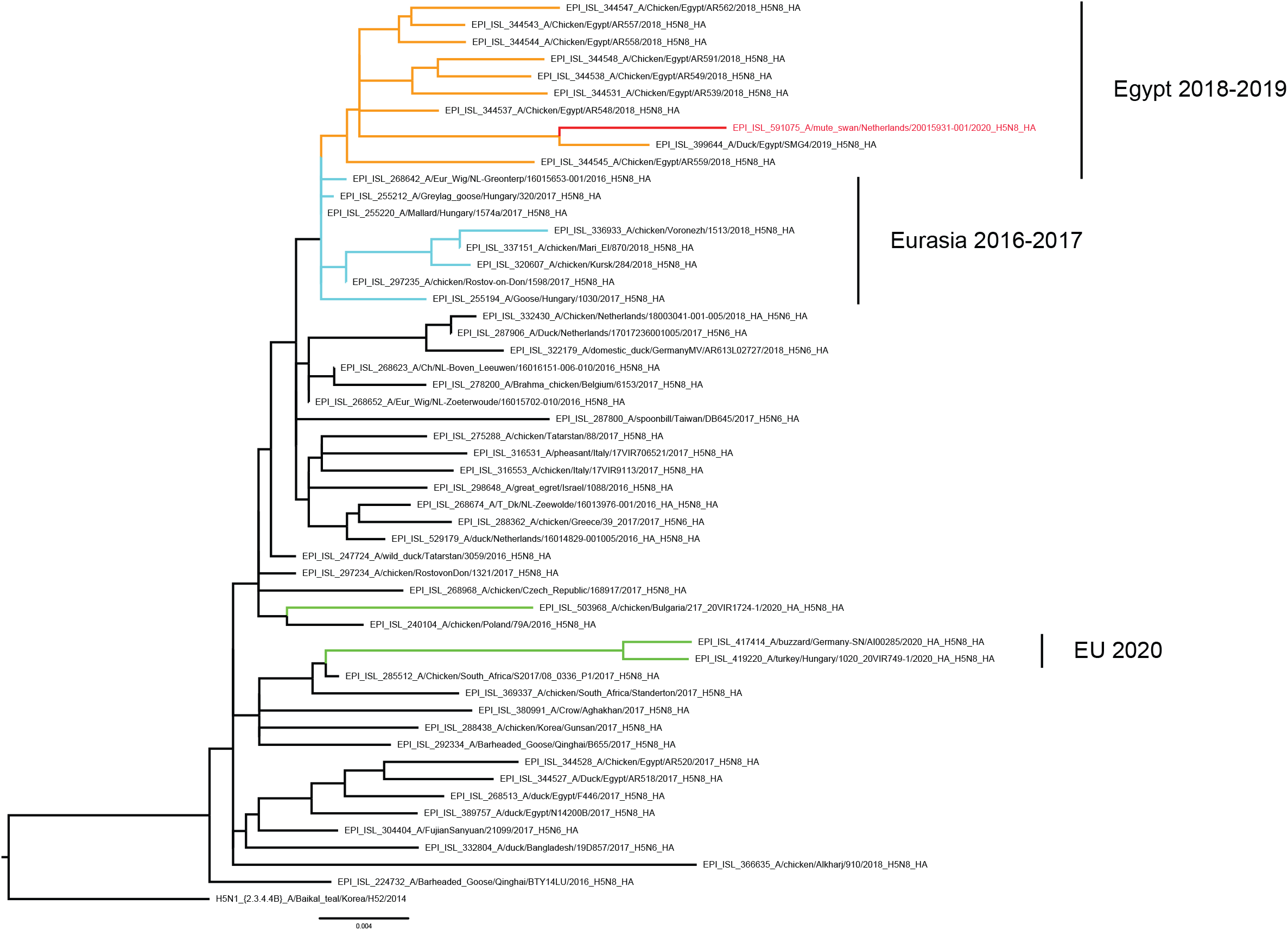
Phylogenetic analysis was performed for the HA segment. The optimal phylogenetic tree generated using the maximum likelihood method (RAxML v8.2.12 with 100 bootstrap replicates) is shown and drawn to scale. The GISAID accession numbers of the viruses are shown in the trees. The H5N8 virus isolated in the Netherlands in 2020 is marked in red, the H5N8 viruses isolated in Eastern-Europe, Germany and Bulgaria in 2020 are marked in green.

The genetic distance between the novel H5N8 virus and related H5N8 viruses detected in Egypt and Eurasia appears relatively large, as is demonstrated by the long branch lengths in the phylogenetic trees (Fig S1). To estimate the time to the most recent common ancestor (tMRCA), we performed molecular dating using the Bayesian Skyline coalescent model (BEAST v 1.10.2 software) (Supplementary FigS2). For the H5 segment (Fig 2), a common ancestor of the novel H5N8 virus and the virus detected in Egypt in 2019 (EPI399644) was dated around July 2018 (Fig 2, [node 1]), and with the cluster of viruses from Egypt around March 2017 (Fig 2, [node 2]). The novel H5N8 virus shares a common ancestor with the Eurasian viruses detected in 2016-2017 which was dated to August 2016 (Fig 2, [node 3]). The ancestor of the novel virus and H5N8 viruses detected in Eastern-Europe and Germany in 2020 was predicted in December 2015 (Fig 2, [node 4]). The highest posterior density interval and the posterior values are summarized for all segments in Table 2. Similar dating of ancestral viruses was observed for the other gene segments (FigS2), except MP that shares a common ancestor with H5N8 viruses isolated in Asia and Egypt dated to July 2016 (Fig S2, [node A]).

**Fig 2.**
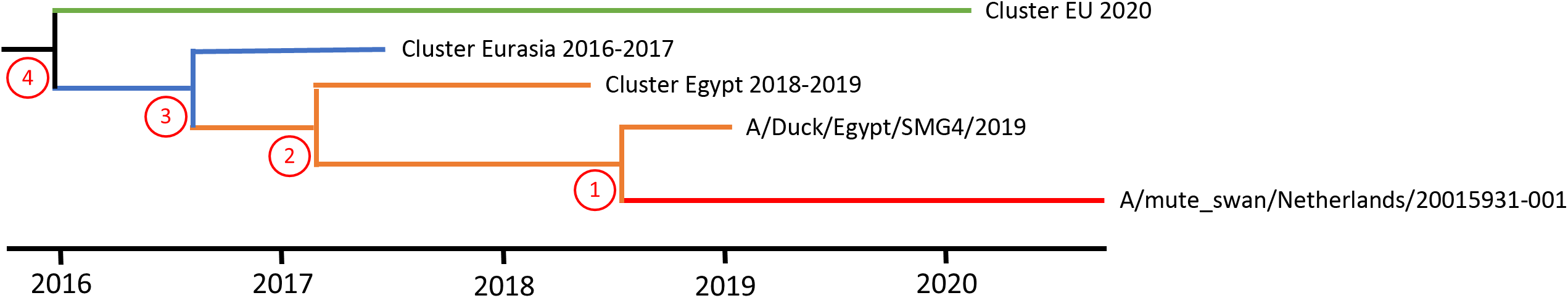
Molecular dating of the HA gene segment. The Bayesian coalescent method was used to estimate the time to the most recent common ancestor (tMRCA) of the novel H5N8 virus. The schematic representation shows the novel H5N8 virus in red, viruses detected in Egypt 2017-2018 and 2019 in orange, Eurasian viruses in blue and the viruses from Eastern-Europe and Germany in 2020 in green.

**Table 2:**
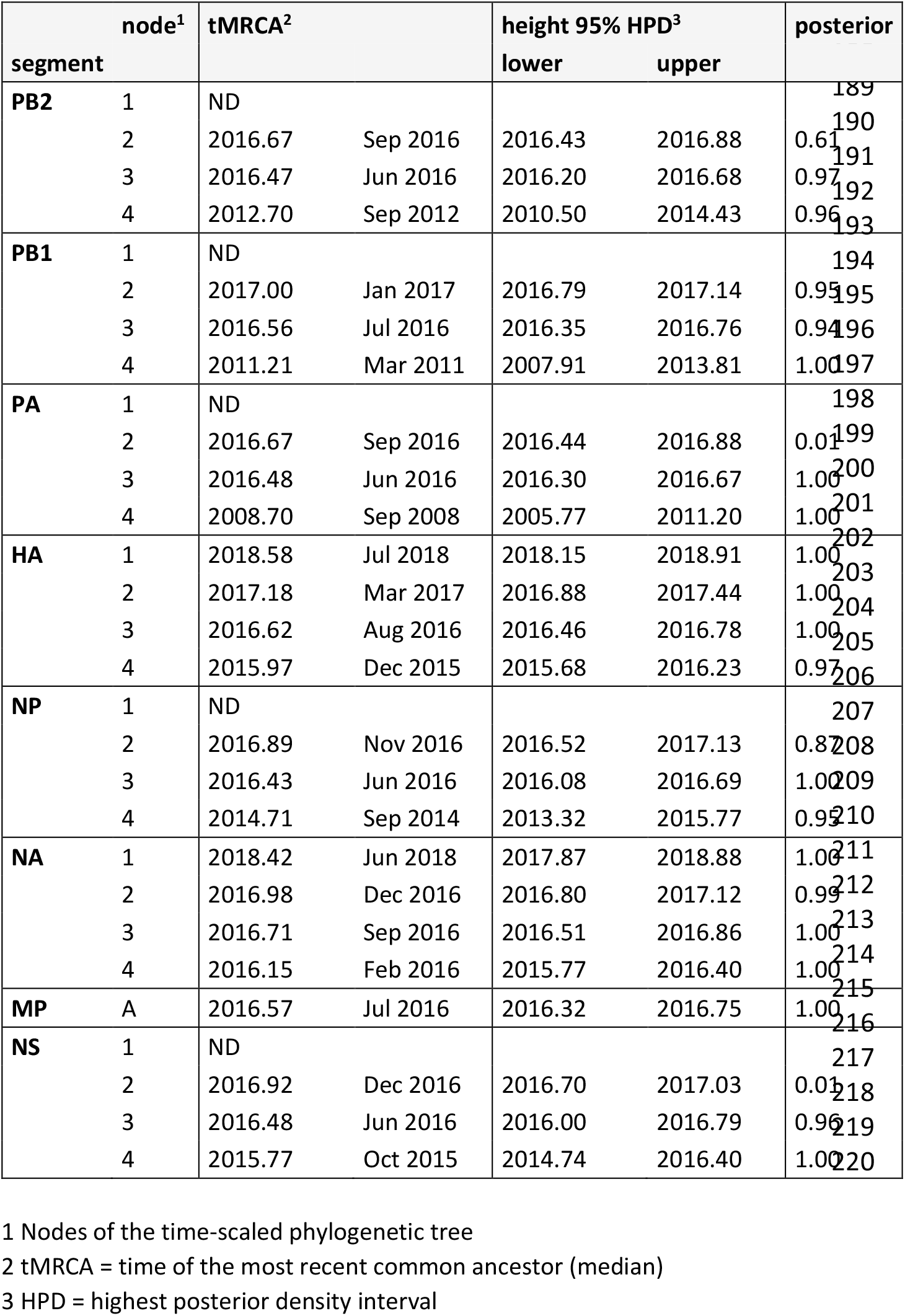
calculated tMRCA, with 95% credible interval and posterior value

The molecular dating analysis suggests that the ancestor of the novel H5N8 virus circulated in this genetic constitution since March 2017, causing outbreaks in Egypt in 2018-2019. Thereafter, the virus remained undetected until its incursion into the Netherlands in October 2020. The virus incursion is not related to outbreaks in Eastern-Europe, Germany, and Bulgaria earlier in 2020. Furthermore, as related viruses were detected in Egypt in 2018-2019, it seems unlikely that the H5N8 virus remained circulating undetected in the Netherlands since 2016. As mute swans do not migrate over long distances, the virus may have been introduced by other waterfowl species of the orders Anseriformes and Charadriiformes migrating to between breeding grounds in North-Russia and wintering sites in the Netherlands. No HPAI H5N8 viruses or wild bird mortality was reported from this (breeding) region proceeding the migration season. However, starting in September 2020, several reports of HPAI H5N8 viruses in wild birds and poultry in the South-Russia and North-Kazakhstan were reported by the responsible national authorities (OIE; https://www.oie.int/en/animal-health-in-the-world/update-on-avian-influenza/2020/). Some waterfowl species, such as Eurasian wigeon *(Anas penelope)*, Tufted duck *(Aythya fuligula)* and White-fronted goose *(Anser albifr*ons), are known to reside in these regions of Russia and Kazakhstan and migrate to the Netherlands for wintering (SOVON; http://vogeltrekatlas.nl/soortzoek2.html). The detection of HPAI H5N8 virus in a Eurasian wigeon found dead near the location of the dead swans suggests that this bird species may have introduced the virus in the Netherlands. As the sequences of the viruses detected in Russia and Kazakhstan are currently unknown, the relationship between these viruses and the virus detected in the Netherlands remains to be determined.

## Conclusions

A novel HPAI H5N8 virus was detected in the Netherlands in October 2020 in mute swans found dead. The virus is not closely related to H5N8 viruses causing outbreaks in Eastern-Europe, Germany and Bulgaria in the first half year of 2020, but shares a putative common ancestor with viruses last detected in Egypt in 2018-2019, which dates around March 2017. The virus was likely introduced by waterfowl migrating to wintering sites in the Netherlands. In October, wild bird migration is ongoing and millions of wild birds will reach their wintering sites in Europe in the coming months. This early detection of HPAI H5N8 virus in the Netherlands predicts a high risk for the poultry industry in Europe for the 2020-2021 winters season.

## Acknowledgments

We thank Alex Bossers for excellent NGS facilities within WBVR. This work was funded by the Dutch Ministry of Agriculture, Nature and Food Quality (project WOT-01-003-087 and KB-37-003-015). We acknowledge the authors and submitting laboratories of the sequences from the GISAID EpiFlu Database (detailed in supplementary file S3).

## Author Bio

Dr Beerens is a senior scientist and head of the National Reference Laboratory for Avian Influenza and Newcastle Disease in the Netherlands. Her research interests focus on molecular virology, genetics, and virus evolution.

## Address for correspondence

Nancy Beerens, Wageningen Bioveterinary Research, Division of Virology, P.O. Box 65, 8200 AB, Lelystad, The Netherlands. Phone: +31 320 238217

## Appendix: Supplementary Figure legends

Fig S1: Phylogenetic analysis was performed for each gene segment. Related sequences were obtained from GISAID’s EpiFlu database (http://www.gisaid.org)(9). For HA additional sequences form H5 clade 2.3.4.4b were collected from the EpiFlu database and clustered using the CD-HIT algoritm (10), using identity setting of 0.985. Cluster representatives were used in the analysis of HA in addition to the related sequences from the GISAID blast analyses. Sequences were aligned using MAFFT v7.427 (11). Maximum likelihood trees based on the general time reversible model with a gamma-distributed variation of rates and 100 bootstraps were generated using RA×ML v8.2.12 (12). The GISAID accession numbers of the viruses are shown in the trees (details see table S3). The H5N8 virus isolated in the Netherlands in 2020 is marked in red, the H5N8 viruses isolated in Eastern-Europe, Germany and Bulgaria in 2020 are marked in green.

Fig S2: Molecular dating was performed for the all gene segments. The datasets of the ML tree analysis (see Fig S1) were used for time scaled phylogenies, reconstructed using a Bayesian Markov chain Monte Carlo (MCMC) framework implemented in the BEAST software package (v 1.10.2; (13). The analysis was carried out with the SRD06 nucleotide substitution model, the Bayesian Skyline coalescent model and an uncorrelated lognormal relaxed molecular clock and MCMC runs of 1×10^8^ states sampling each 1×10^4^ steps were run to obtain an effective sample size of >200. Maximum clade credibility (MCC) trees were reconstructed with 10% burn-in and the posterior distribution of relevant parameters were assessed in FigTree (v 1.4.4). The tMRCA for the numbered nodes is listed in Table 2, as is the credible interval and posterior value. GISAID accession numbers are shown in the trees (for details see Supplementary file S3). The H5N8 virus isolated in the Netherlands in 2020 is marked in red, the H5N8 viruses isolated in Eastern-Europe, Germany and Bulgaria in 2020 are marked in green.

