## Supplementary figures and images for "Novel incursion of a highly pathogenic avian influenza subtype H5N8 virus in the Netherlands, October 2020"

### Suppl. Fig S1

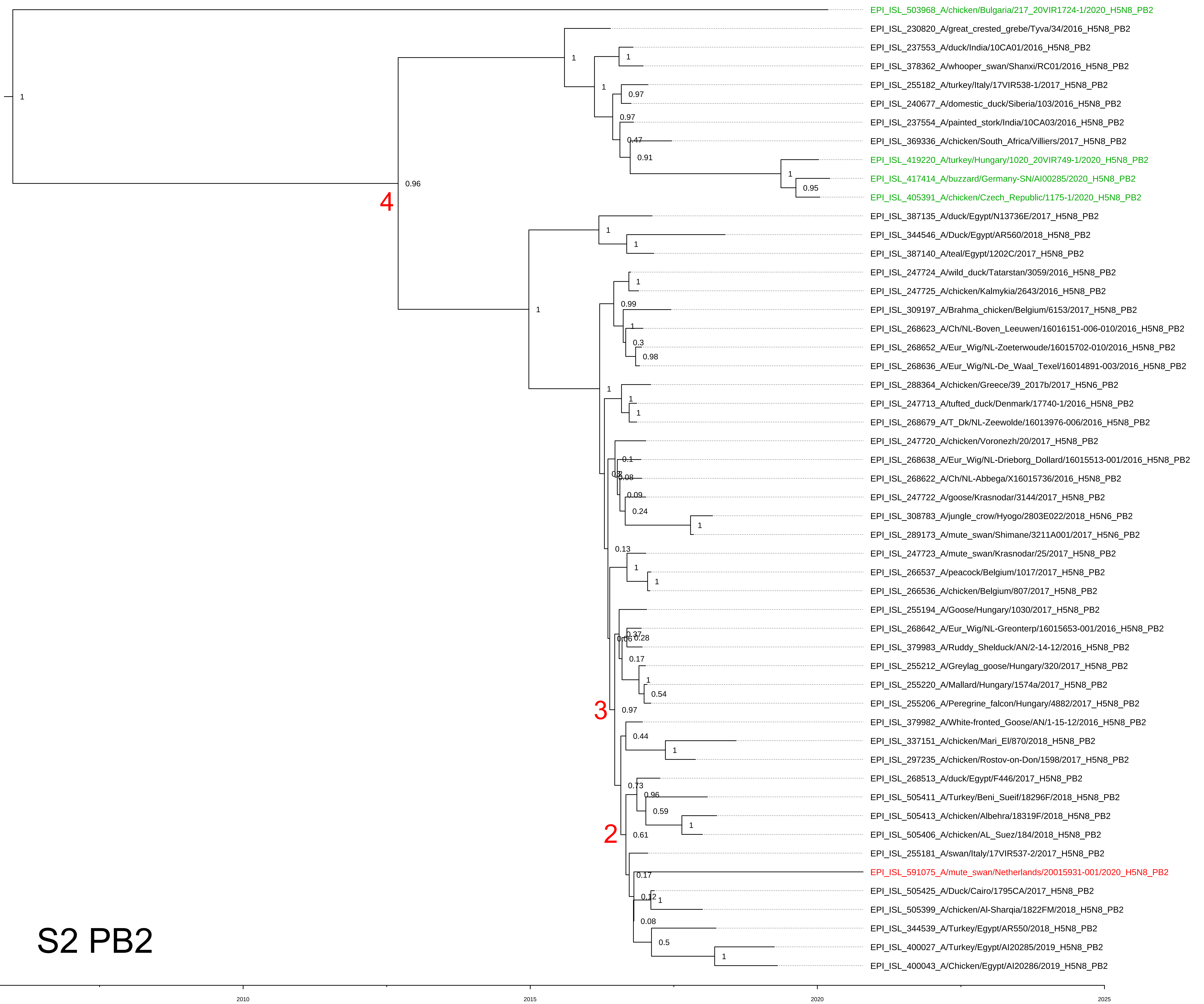

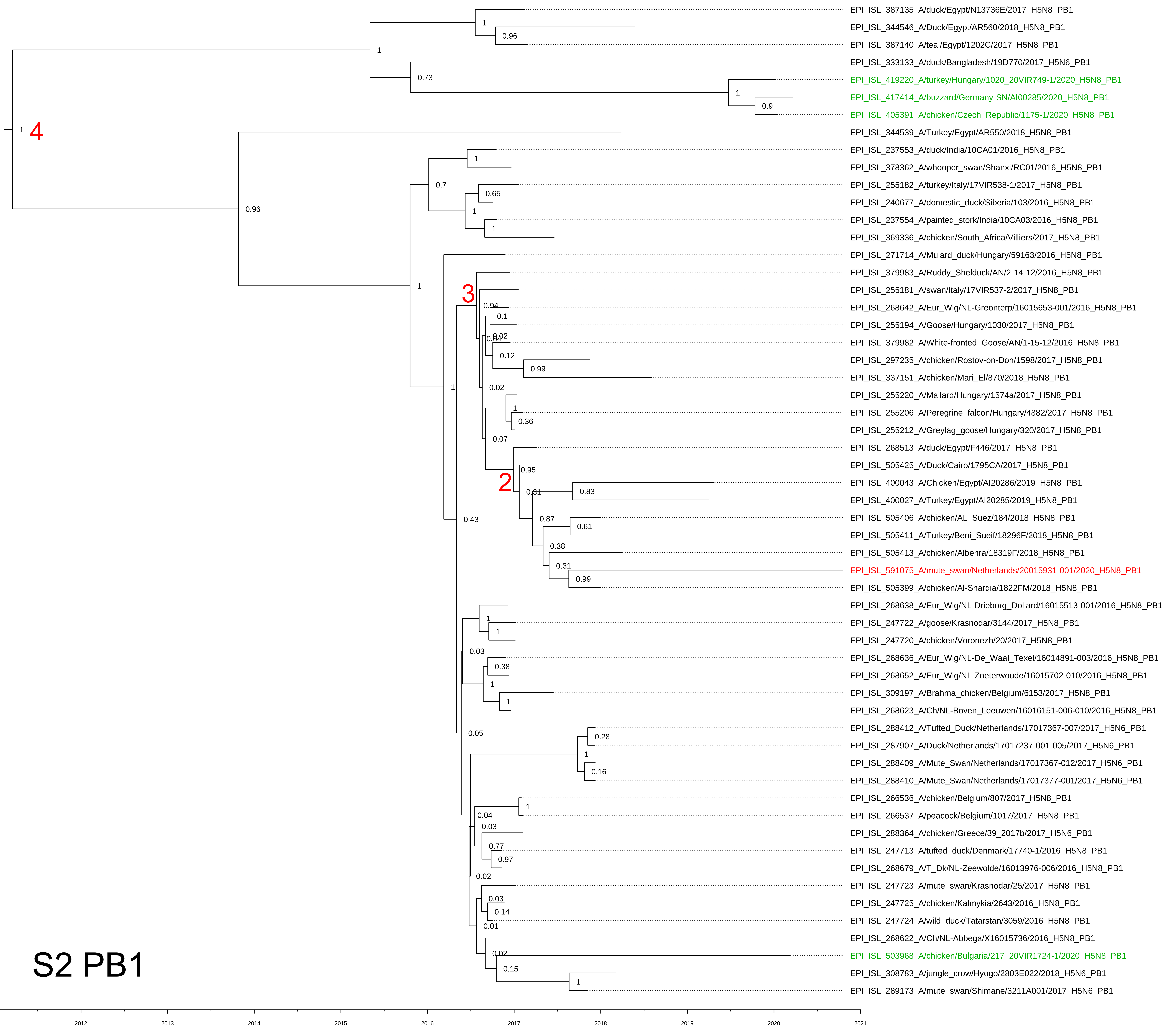

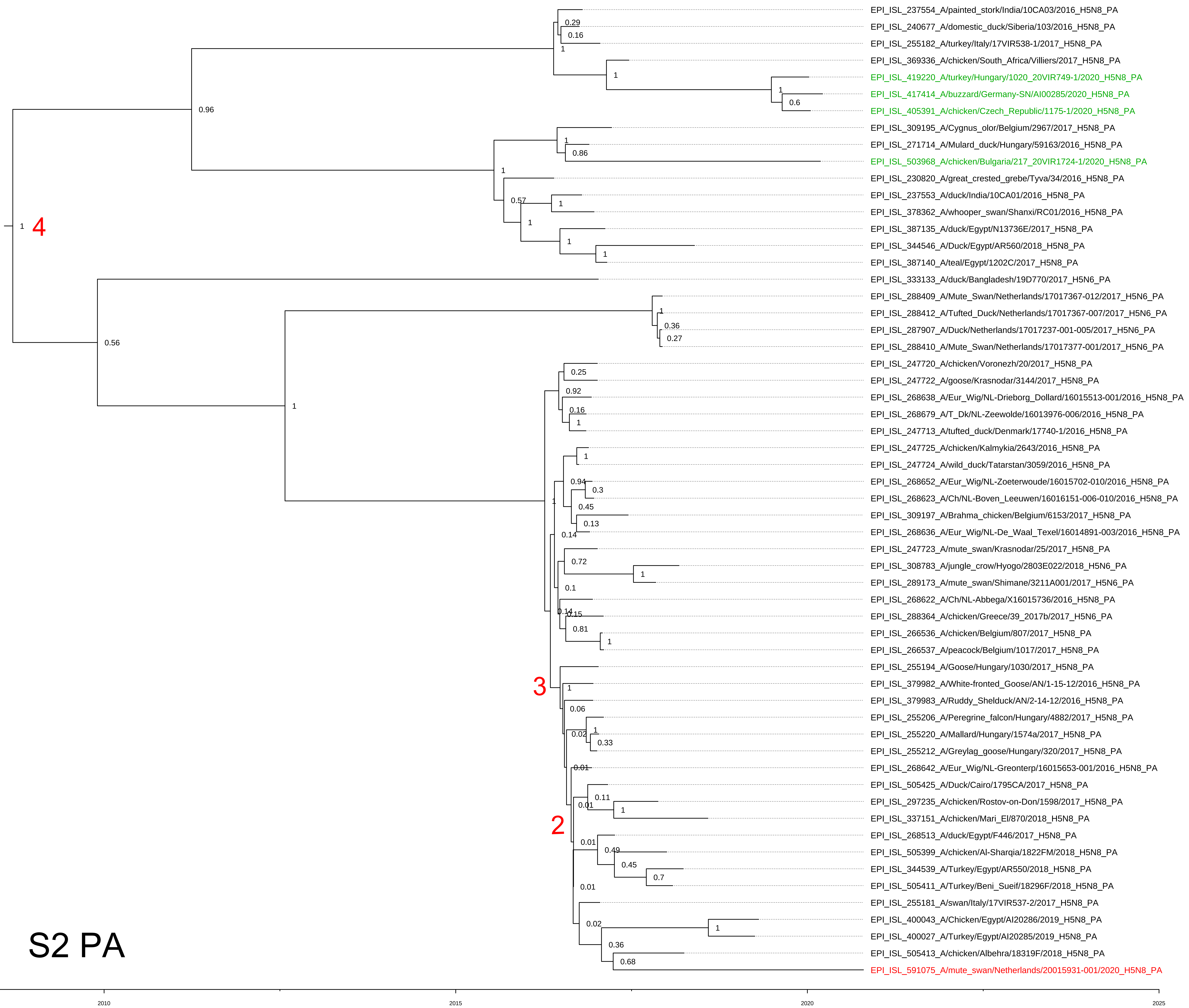

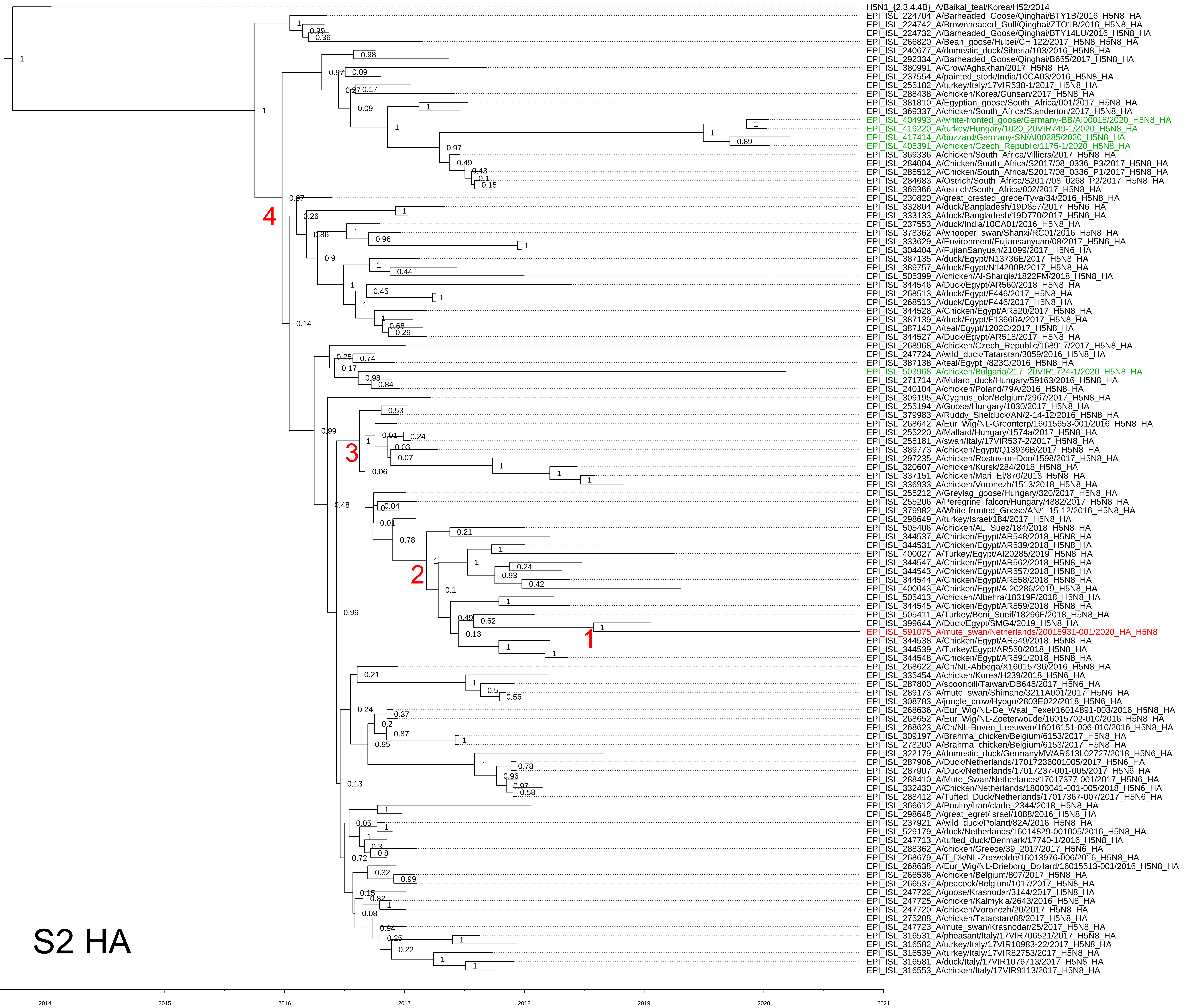

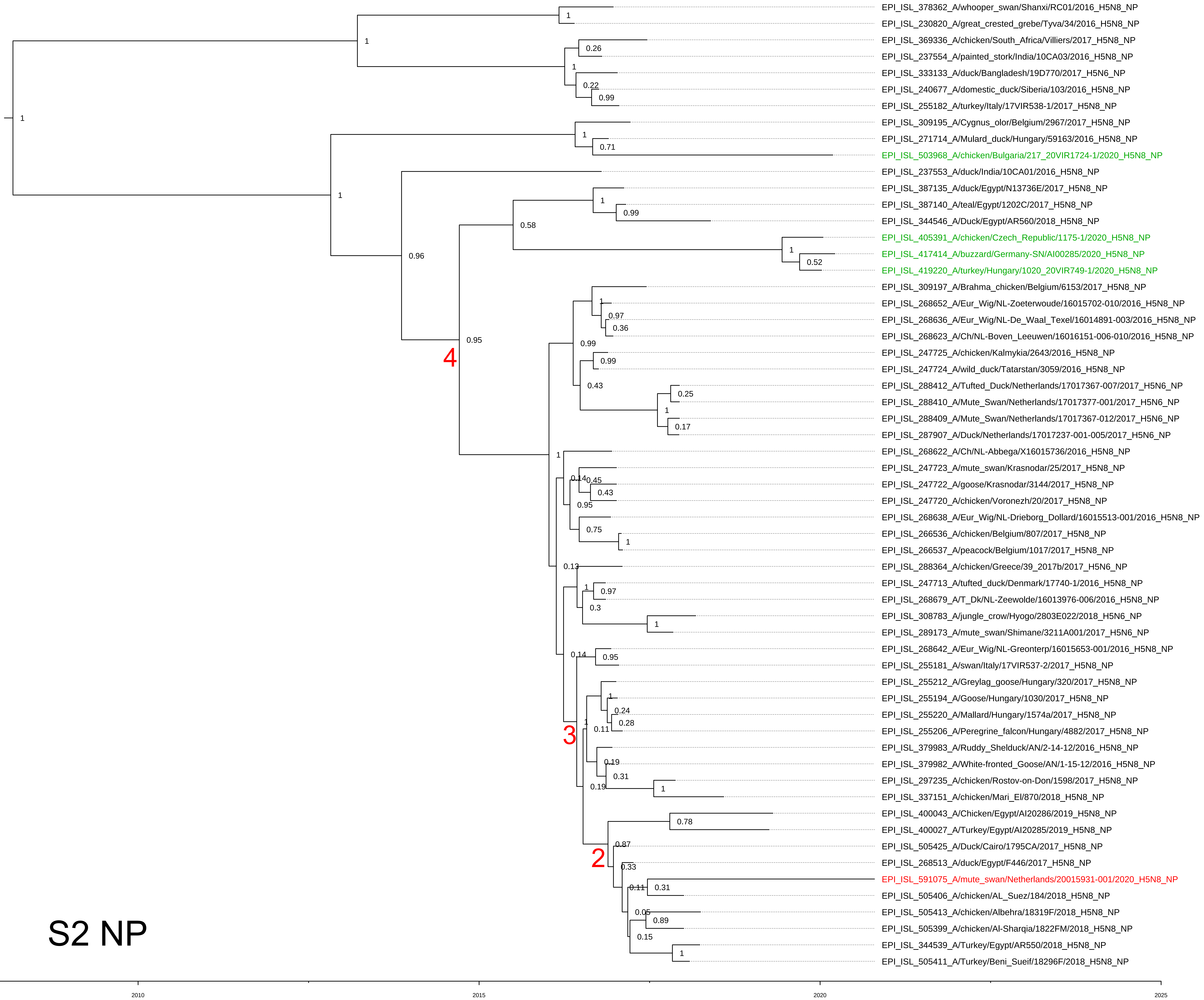

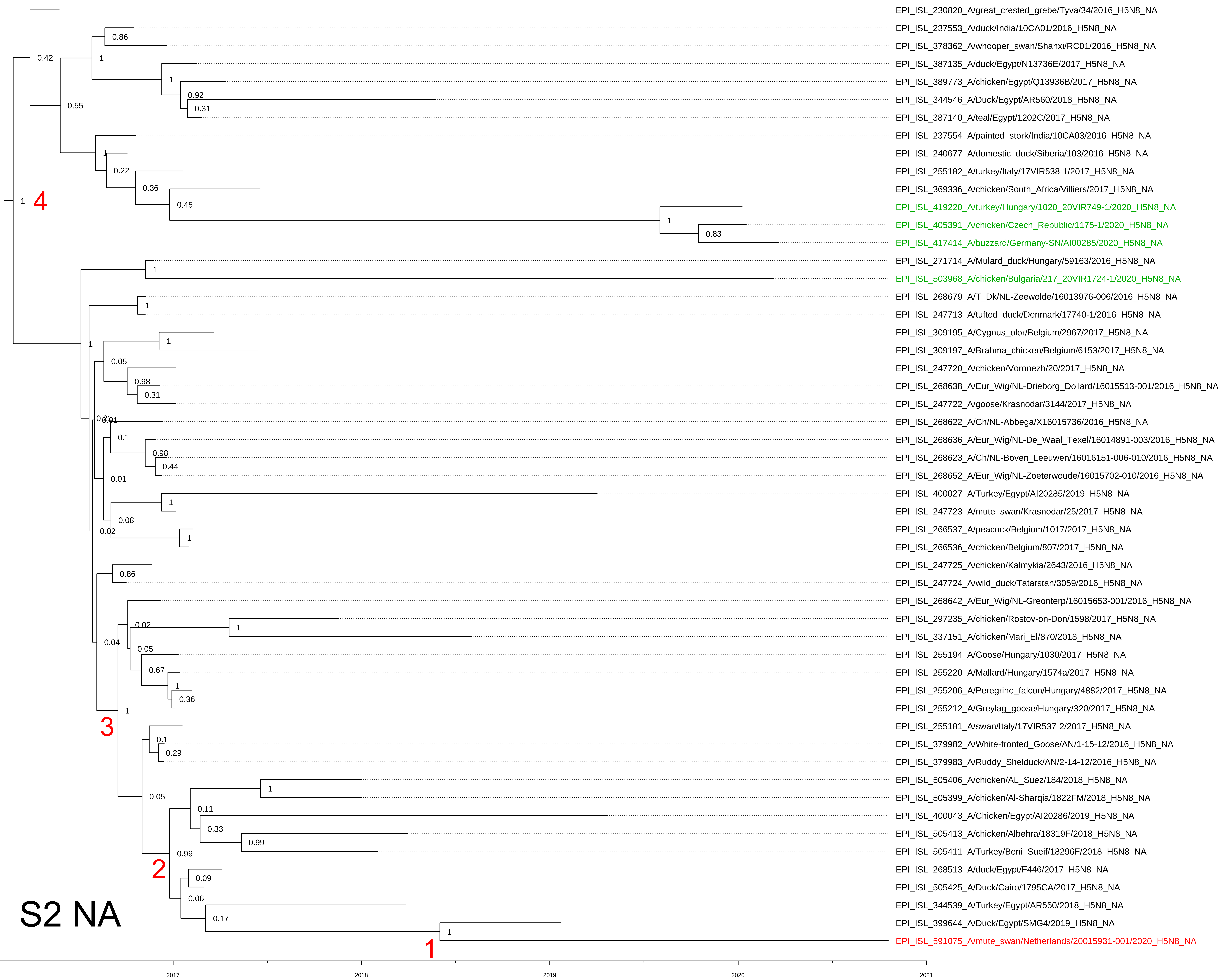

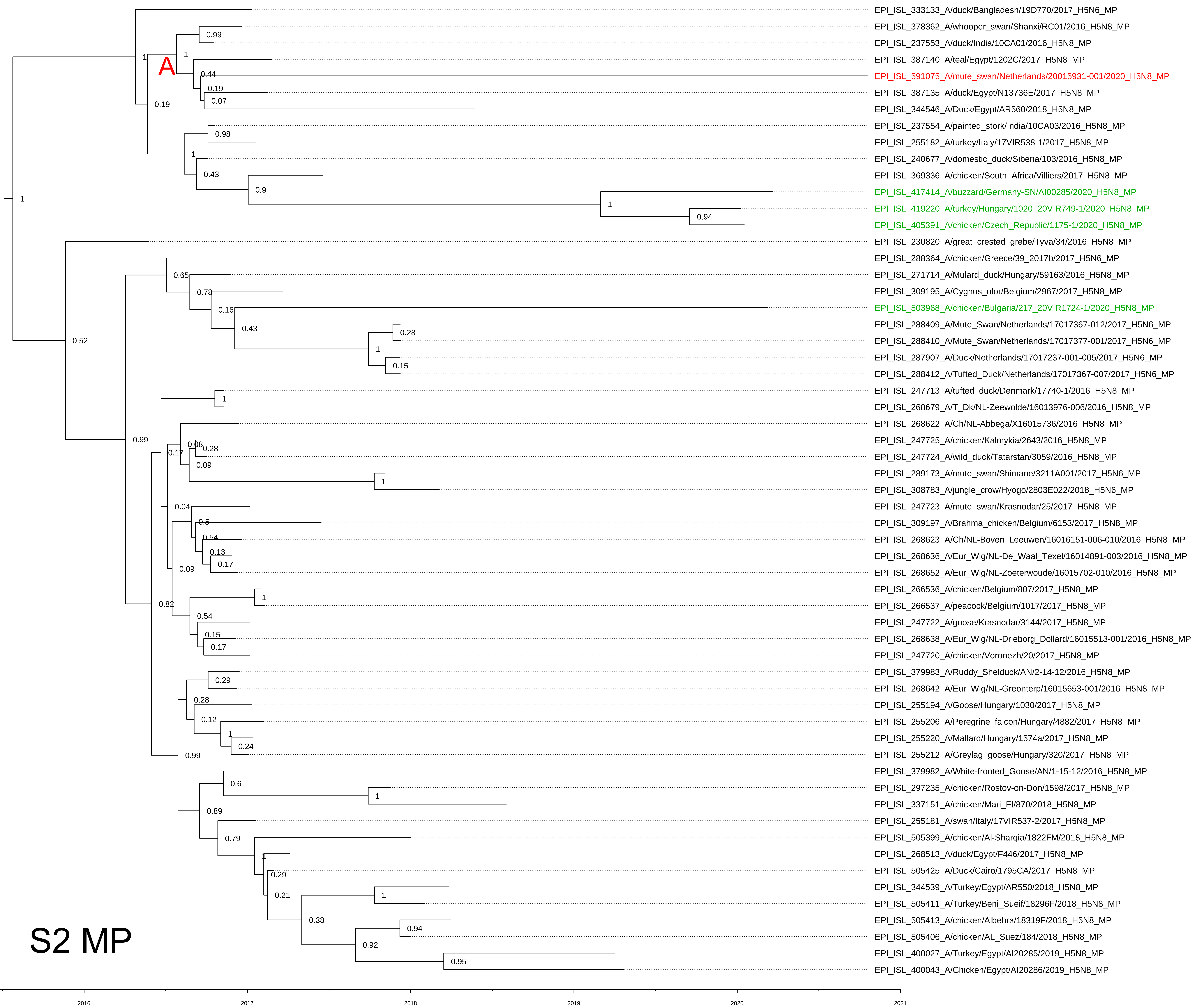

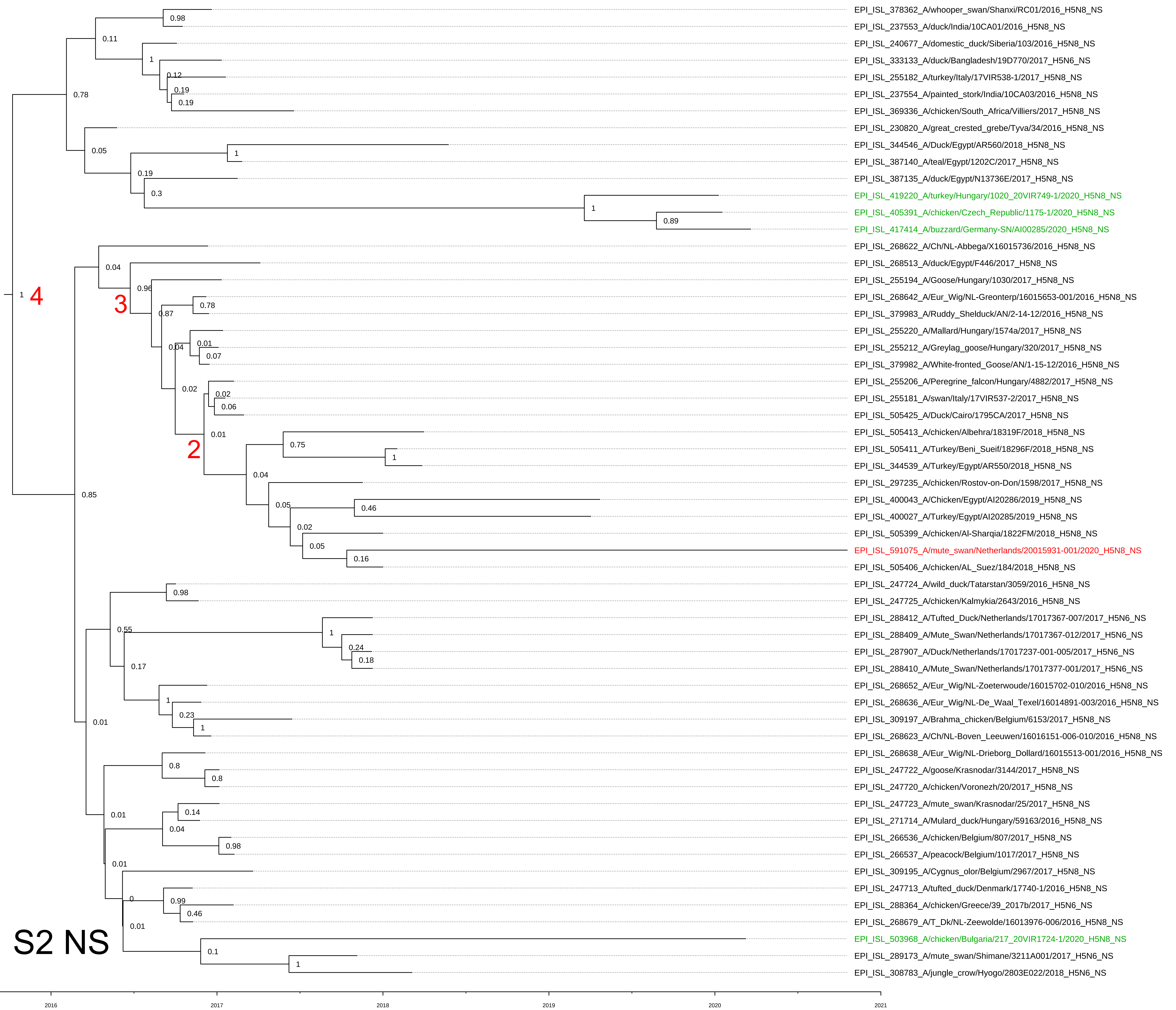

### Suppl. Fig S2

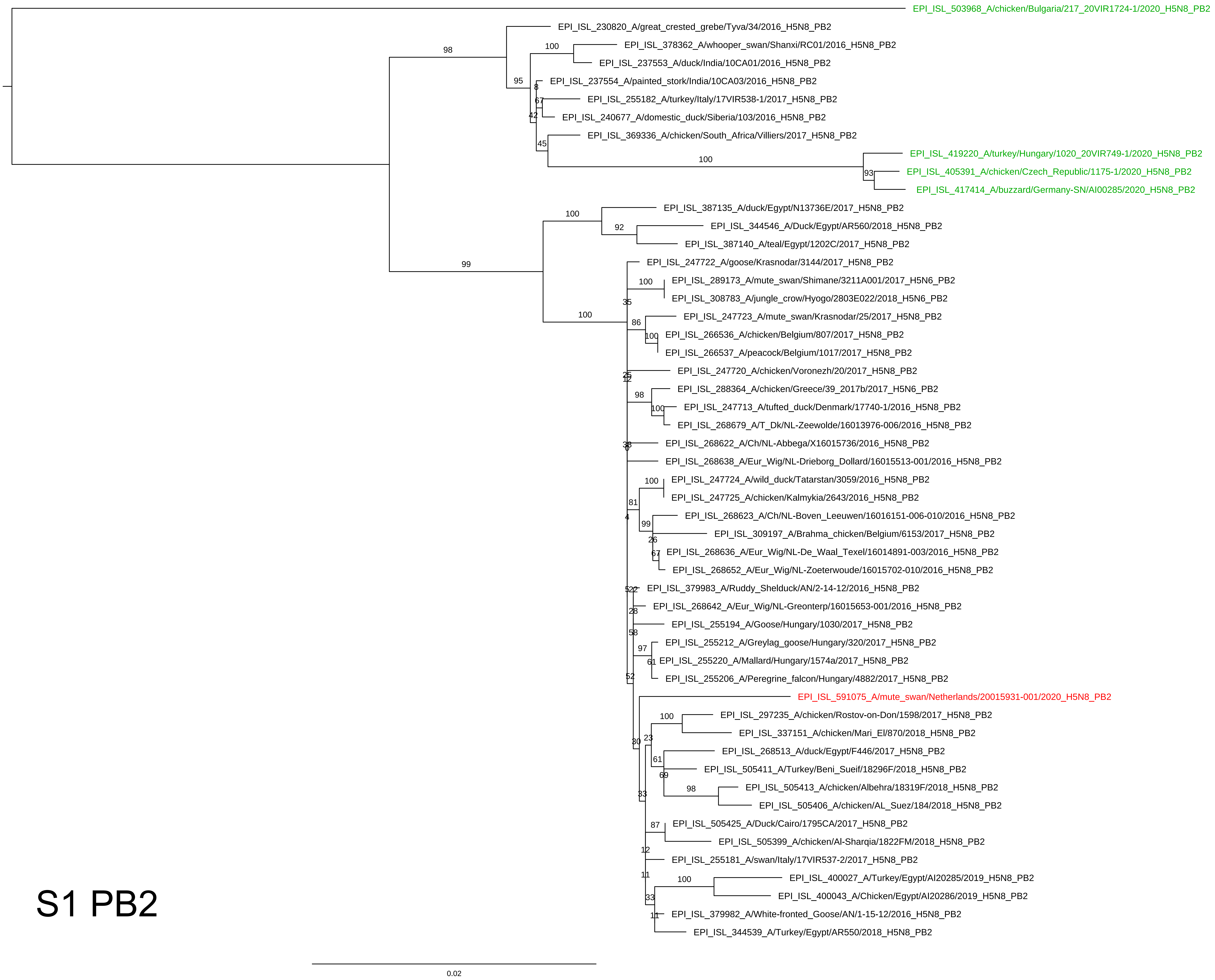

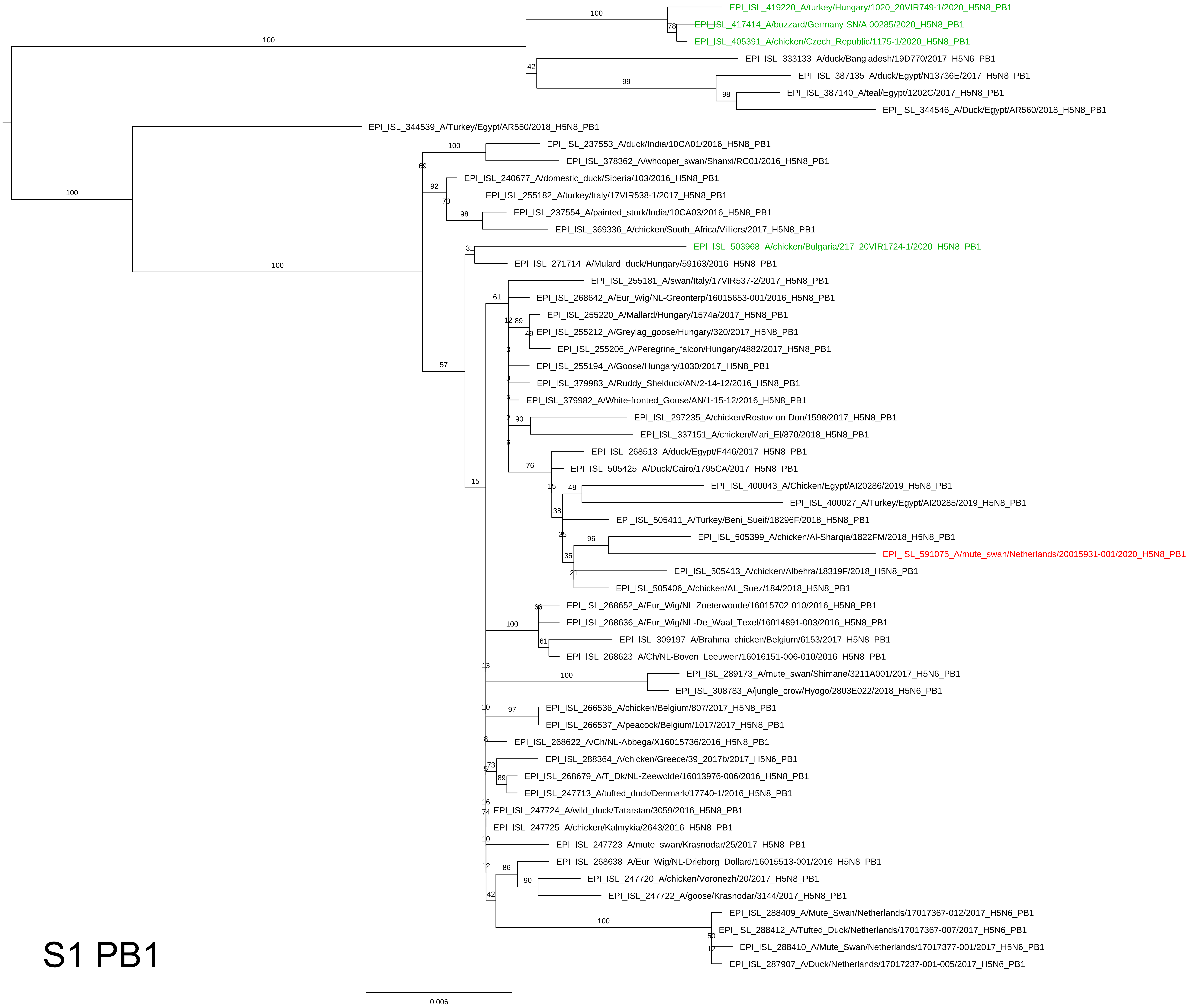

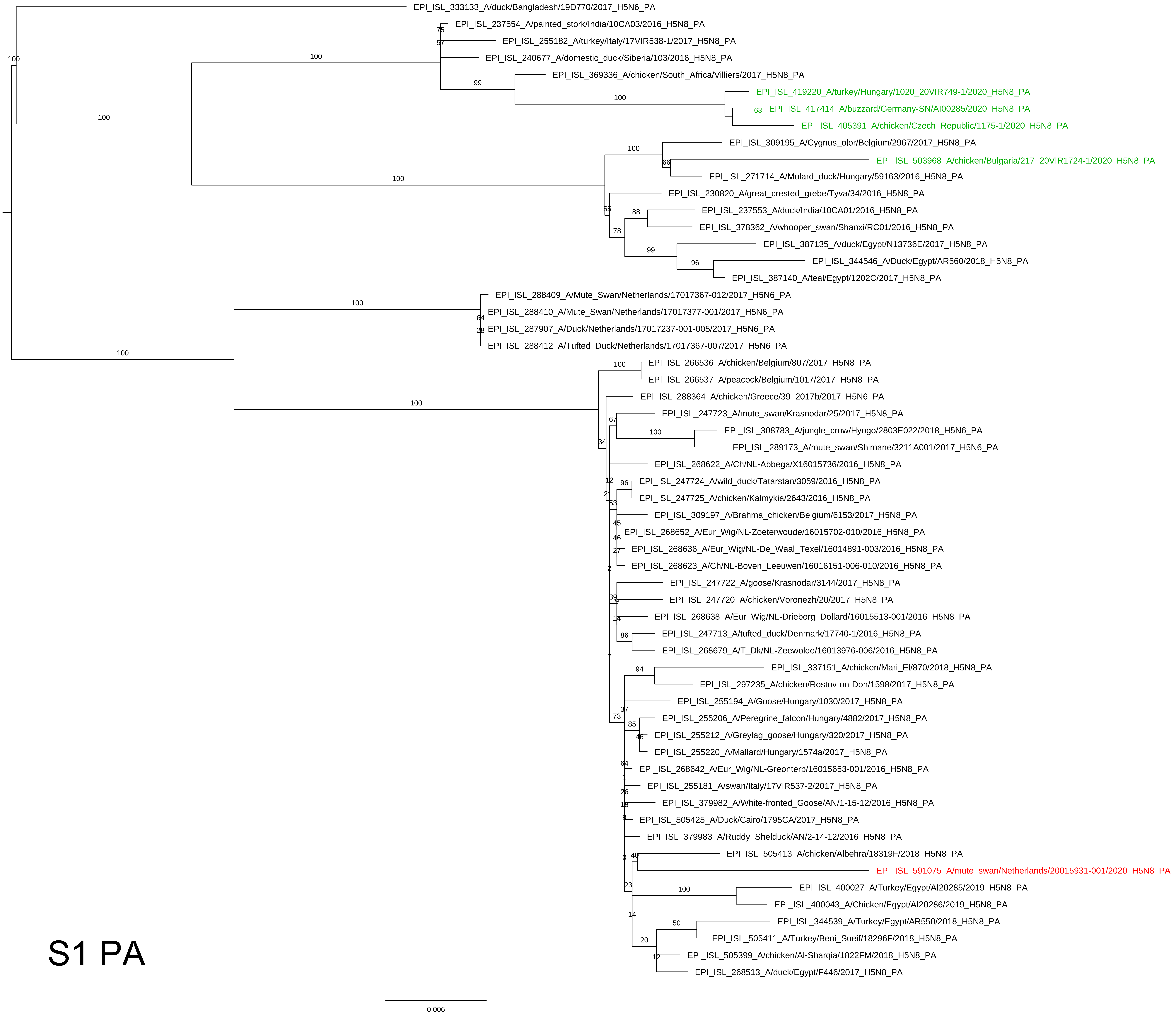

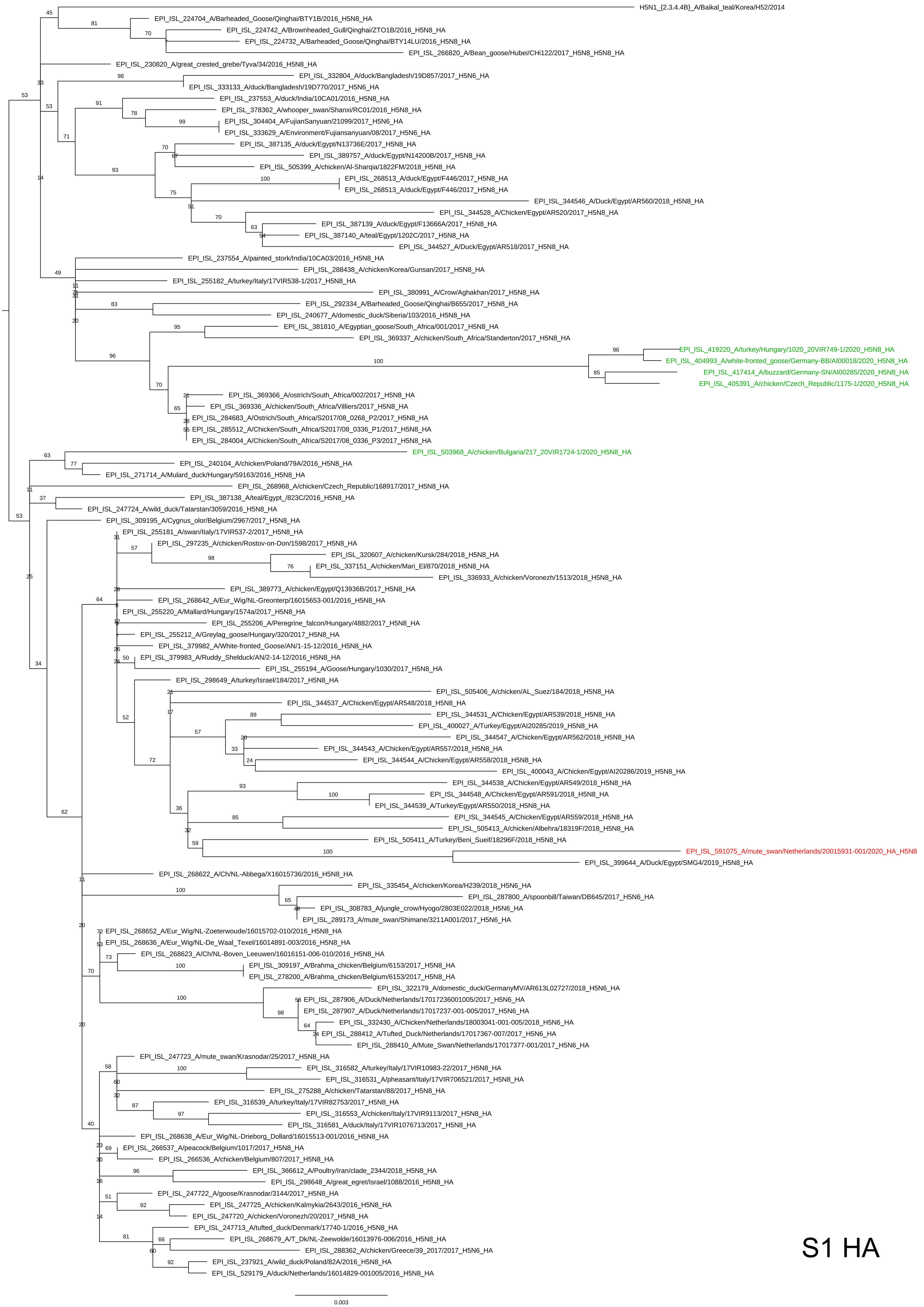

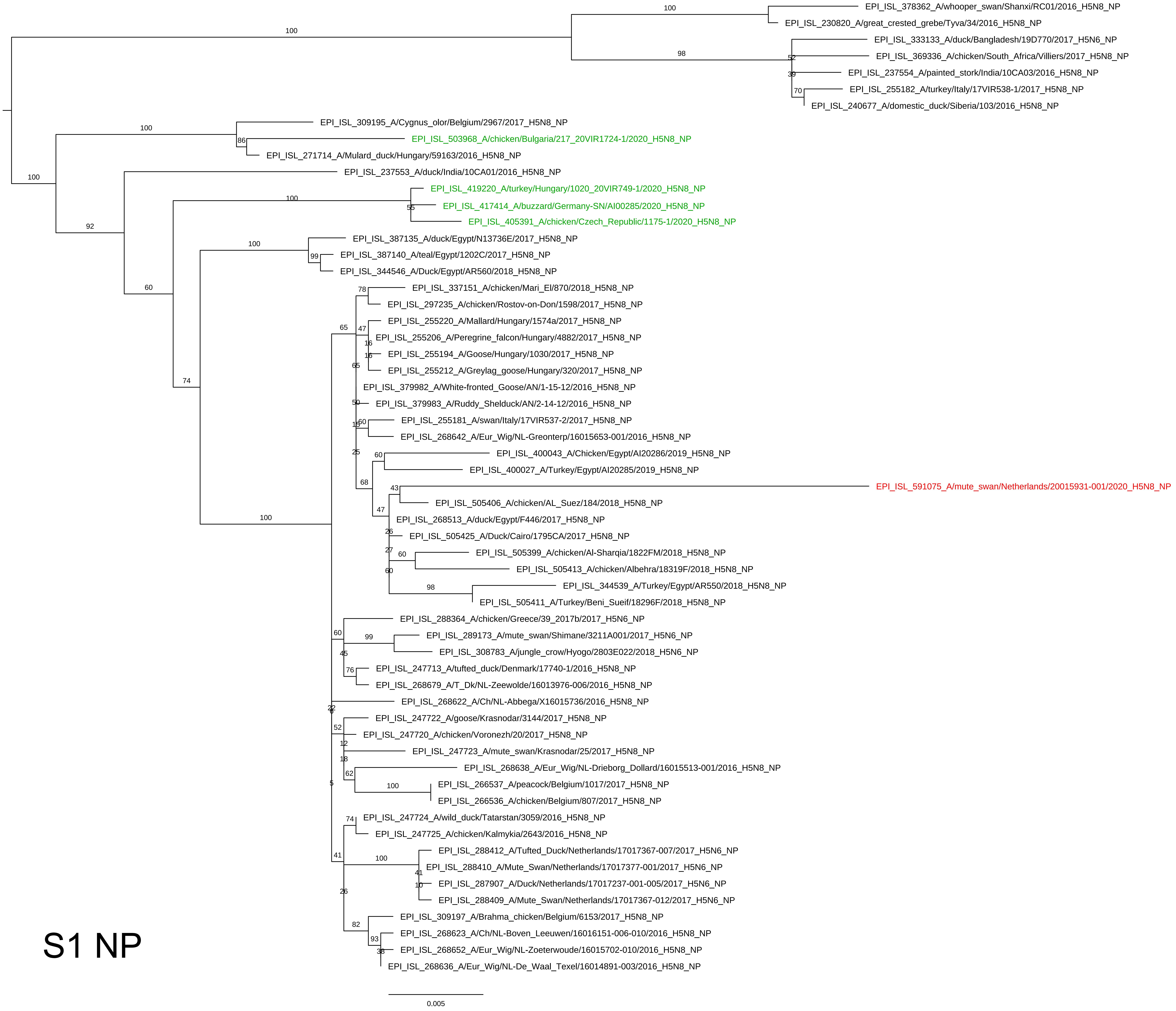

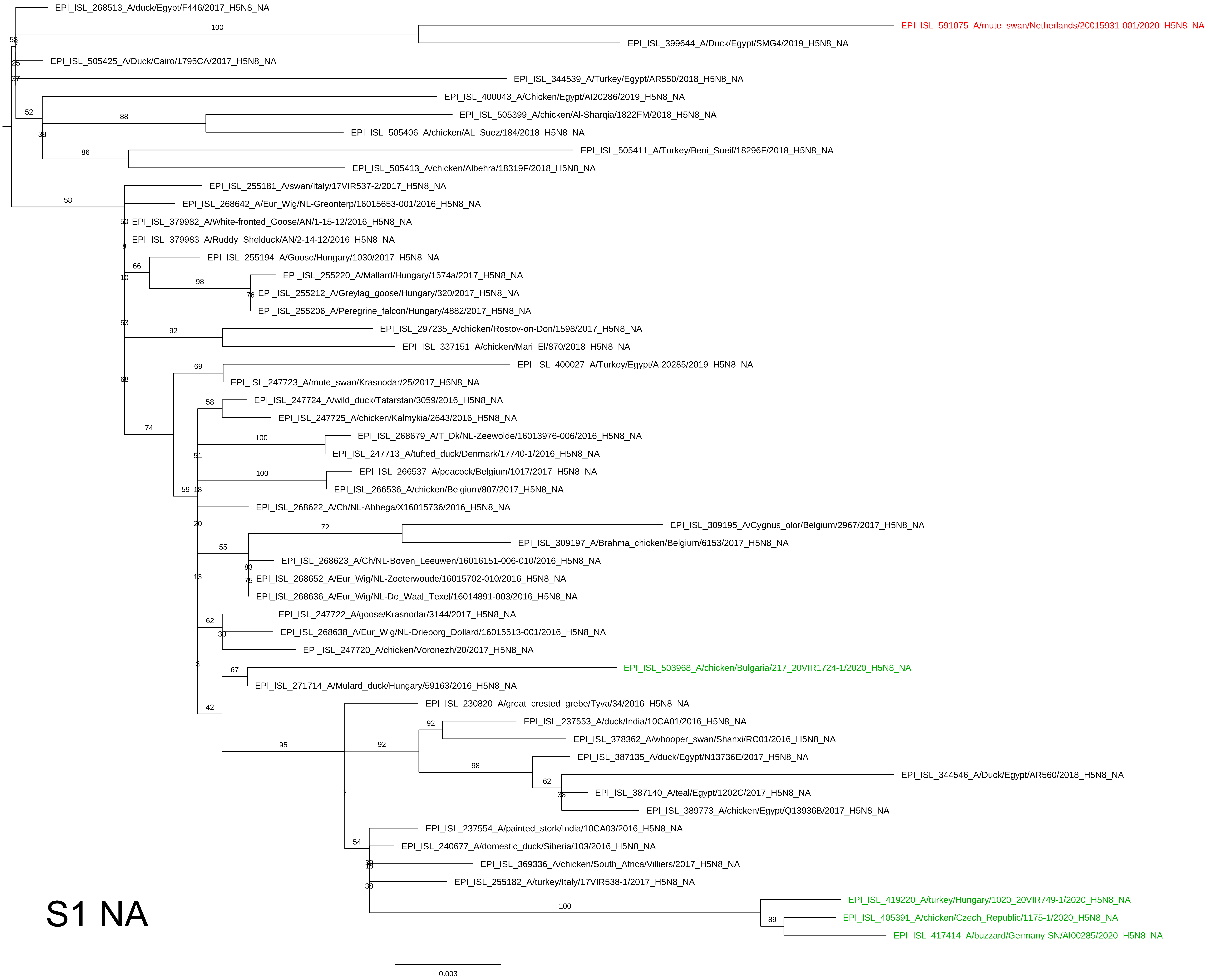

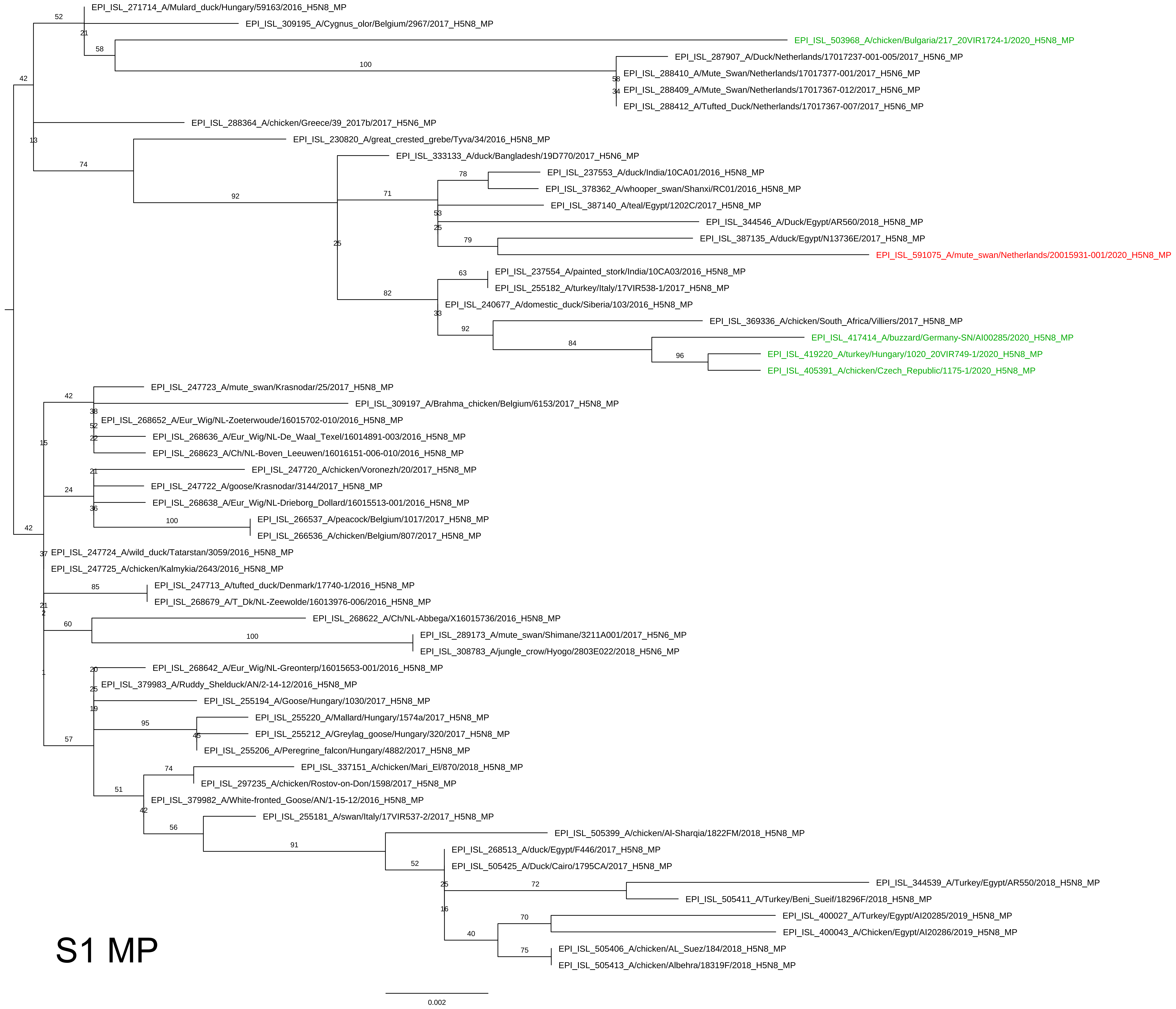

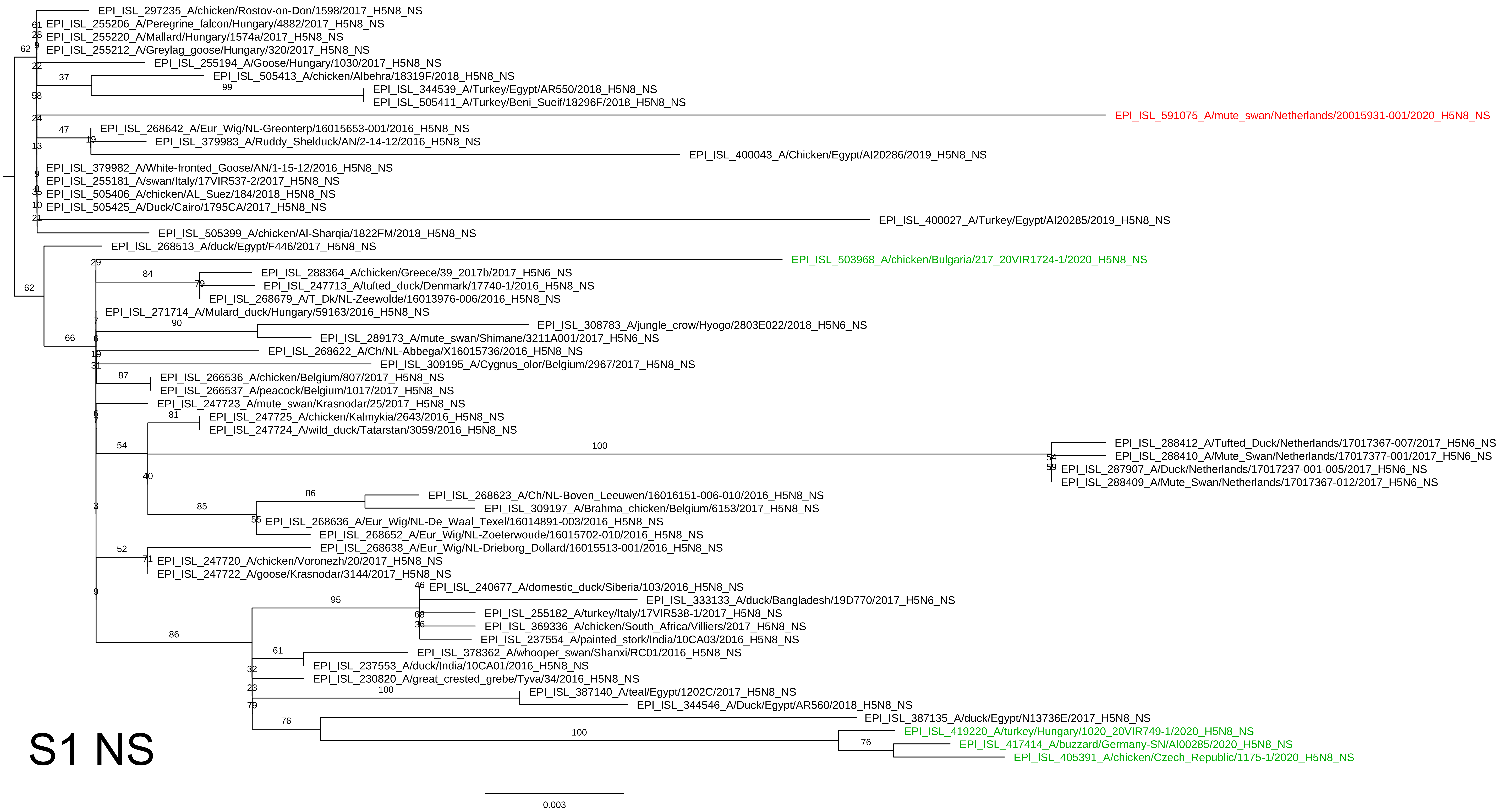
