## Supplementary material for "Novel incursion of a highly pathogenic avian influenza subtype H5N8 virus in the Netherlands, October 2020": Suppl. FileS3

We gratefully acknowledge the authors, originating and submitting laboratories of the sequences from GISAID's EpiFlu™ Database on which this research is based. The list is detailed below.
All submitters of data may be contacted directly via www.gisaid.org

| Isolate_Id | PB2 Segment | PB1 Segment_I | PA Segment_Id | HA Segment_Id | NP Segment_Id | NA Segment_Id | MP Segment_Id | NS Segment_Id | Isolate_Name | Subtype | Passage_His | Location | Host | Isolate_Submitter | Submitting_Lab | Submitting_S | Authors | Originating_Lab | Originating_S | Collection_D | Update_Date | Submission_ |
|---|---|---|---|---|---|---|---|---|---|---|---|---|---|---|---|---|---|---|---|---|---|---|
| EPI_ISL_224704 | EPI774110\|A/E | EPI774111\|A/Ba | EPI774112\|A/Bar- | EPI774113\|A/Bar- | EPI774114\|A/Bar- | EPI774115\|A/Bar- | EPI774116\|A/Bar- | EPI774117\|A/Bar- | A/Bar-headed Goose/Qinghai/BTY1-B/2016 | A / H5N8 | egg1 | Asia / China / | Anser indicus | Jianjun Chen (Wuhan Institute of Virology / Chinese Academy of Sciences) | Wuhan Institute of Virology | | | State Key Laboratory of Virology and Wuhan Institute of | | 2016-05-09 | 2016-07-12 | 2016-07-12 |
| EPI_ISL_224732 | EPI774341\|A/E | EPI774342\|A/Ba | EPI774343\|A/Bar- | EPI774344\|A/Bar- | EPI774345\|A/Bar- | EPI774346\|A/Bar- | EPI774347\|A/Bar- | EPI774348\|A/Bar- | A/Bar-headed Goose/Qinghai/BTY14-LU/2016 | A / H5N8 | egg1 | Asia / China / | Anser indicus | Jianjun Chen (Wuhan Institute of Virology / Chinese Academy of Sciences) | Wuhan Institute of Virology | | | State Key Laboratory of Virology and Wuhan Institute of | | 2016-05-14 | 2016-07-12 | 2016-07-12 |
| EPI_ISL_224742 | EPI774423\|A/E | EPI774424\|A/Br | EPI774425\|A/Brov | EPI774426\|A/Brov | EPI774427\|A/Brov | EPI774428\|A/Brov | EPI774429\|A/Brov | EPI774430\|A/Brov | A/Brown-headed Gull/Qinghai/ZTO1-B/2016 | A / H5N8 | egg1 | Asia / China / | Larus brunnice | Jianjun Chen (Wuhan Institute of Virology / Chinese Academy of Sciences) | Wuhan Institute of Virology | | | State Key Laboratory of Virology and Wuhan Institute of | | 2016-05-01 | 2016-07-12 | 2016-07-12 |
| EPI_ISL_230820 | EPI823457\|A/g | EPI823458\|A/gre | EPI823459\|A/grea | EPI823460\|A/grea | EPI823461\|A/grea | EPI823462\|A/grea | EPI823463\|A/grea | EPI823464\|A/grea | A/great crested grebe/Tyva/34/2016 | A / H5N8 | | Europe / Russ | Other avian | Artem Fadeev (WHO National Influenza Centre Russian Federation) | WHO National Influenza Centre Russian Federation | | Fadeev,Artem; Komissarov,Andrey; Egorova,Anna; Sintsova, | State Research Center of Virology and Biotechnology (V | | 2016-05-25 | 2016-08-13 | 2016-08-08 |
| EPI_ISL_237553 | EPI858833\|A/d | EPI858834\|A/du | EPI858835\|A/duck | EPI858836\|A/duck | EPI858837\|A/duck | EPI858838\|A/duck | EPI858840\|A/duck | EPI858839\|A/duck | A/duck/India/10CA01/2016 | A / H5N8 | c1 | Asia / India / N | Duck | Shanmuga sundaram Nagarajan (ICAR-National Institute of High Security Animal Dis | ICAR-National Institute of High Security Animal Diseases | | Nagarajan, Shanmugasundaram; Tripathi, Sushil; Kumar, Man | ICAR-National Institute of High Security Animal Disease | | 2016-10-17 | 2016-11-10 | 2016-11-10 |
| EPI_ISL_237554 | EPI858841\|A/p | EPI858842\|A/pa | EPI858843\|A/pain | EPI858844\|A/pain | EPI858845\|A/pain | EPI858846\|A/pain | EPI858847\|A/pain | EPI858848\|A/pain | A/painted stork/India/10CA03/2016 | A / H5N8 | c1 | Asia / India / N | Other avian | Shanmuga sundaram Nagarajan (ICAR-National Institute of High Security Animal Dis | ICAR-National Institute of High Security Animal Diseases | | 'Nagarajan, Shanmugasundaram; Shukla, Shweta; Kumar, Mai | ICAR-National Institute of High Security Animal Disease | | 2016-10-20 | 2016-11-10 | 2016-11-10 |
| EPI_ISL_237921 | EPI860233\|A/w | EPI860234\|A/wil | EPI860235\|A/wild | EPI860231\|A/wild | EPI860236\|A/wild | EPI860232\|A/wild | EPI860237\|A/wild | EPI860238\|A/wild | A/wild duck/Poland/82A/2016 | A / H5N8 | original sampl | Europe / Pola | Duck | Edyta Świętoń (National Veterinary Research Institut Poland, PIWet-PIB) | National Veterinary Research Institut Poland, PIWet-PIB | | Świętoń, E.; Śmietanka, K. | | | 2016-11-02 | 2016-11-18 | 2016-11-16 |
| EPI_ISL_240104 | EPI959432\|A/c | EPI959431\|A/chi | EPI959430\|A/chicl | EPI869927\|A/chicl | EPI959428\|A/chicl | EPI869933\|A/chicl | EPI959427\|A/chicl | EPI959429\|A/chicl | A/chicken/Poland/79A/2016 | A / H5N8 | organ sample | Europe / Pola | Gallus gallus | Edyta Świętoń (National Veterinary Research Institut Poland, PIWet-PIB) | National Veterinary Research Institut Poland, PIWet-PIB | | E. Swieton, K. Smietanka | | | 2016-12-17 | 2017-01-02 | 2017-01-02 |
| EPI_ISL_240677 | EPI926610\| A/ | EPI926611\| A/do | EPI926612\| A/don | EPI926613\| A/don | EPI926614\| A/don | EPI926615\| A/don | EPI926616\| A/don | EPI926617\| A/don | A/domestic duck/Siberia/103/2016 | A / H5N8 | Egg 2 | Europe / Russ | Duck | Ivan Sobolev (Research Institute of Experimental and Clinical Medicine) | Research Institute of Experimental and Clinical Medicine | | Sharshov, K.A.; Kurskaya, O.G.; Alexeev, A.Y.; Sobolev, I.A.; | Research Institute of Experimental and Clinical Medicine | | 2016-10-04 | 2019-01-25 | 2017-01-08 |
| EPI_ISL_247713 | EPI909361\|A/t | EPI909362\|A/tuf | EPI909363\|A/tufte | EPI909364\|A/tufte | EPI909365\|A/tufte | EPI909366\|A/tufte | EPI909367\|A/tufte | EPI909368\|A/tufte | A/tufted duck/Denmark/17740-1/2016 | A / H5N8 | egg1 | Europe / Denr | Aythya fuligula | Charlotte Kristiane Hjulsager (Technical University of Denmark) | Animal and Plant Health Agency (APHA) | | Hjulsager,CK; Krog,JS; Larsen,LE; Kvisgaard,LK; Essen,S | Technical University of Denmark | 16-17740-1 | 2016-11-08 | 2017-02-11 | 2017-02-11 |
| EPI_ISL_247720 | EPI909417\|A/c | EPI909418\|A/chi | EPI909419\|A/chicl | EPI909420\|A/chicl | EPI909421\|A/chicl | EPI909422\|A/chicl | EPI909423\|A/chicl | EPI909424\|A/chicl | A/chicken/Voronezh/20/2017 | A / H5N8 | e2 | Europe / Russ | Gallus gallus | Ivan Susloparov (State Research Center of Virology and Biotechnology (VECTOR) | State Research Center of Virology and Biotechnology (VECTOR | | Ivan,Susloparov; Natalya,Goncharova; Natalya,Kolosova; Vasiliy,Marchenko; Alexander,Ryzhikov | | | 2017-01-06 | 2017-02-11 | 2017-02-11 |
| EPI_ISL_247722 | EPI909433\|A/g | EPI909434\|A/go | EPI909435\|A/goos | EPI909436\|A/goos | EPI909437\|A/goos | EPI909438\|A/goos | EPI909439\|A/goos | EPI909440\|A/goos | A/goose/Krasnodar/3144/2017 | A / H5N8 | e2 | Europe / Russ | Anser anser | Ivan Susloparov (State Research Center of Virology and Biotechnology (VECTOR) | State Research Center of Virology and Biotechnology (VECTOR | | Ivan,Susloparov; Natalya,Goncharova; Natalya,Kolosova; Vasiliy,Marchenko; Alexander,Ryzhikov | | | 2017-01-06 | 2017-02-11 | 2017-02-11 |
| EPI_ISL_247723 | EPI909441\|A/m | EPI909442\|A/mu | EPI909443\|A/mute | EPI909444\|A/mute | EPI909445\|A/mute | EPI909446\|A/mute | EPI909447\|A/mute | EPI909448\|A/mute | A/mute swan/Krasnodar/25/2017 | A / H5N8 | e2 | Europe / Russ | Cygnus olor | Ivan Susloparov (State Research Center of Virology and Biotechnology (VECTOR) | State Research Center of Virology and Biotechnology (VECTOR | | Ivan,Susloparov; Natalya,Goncharova; Natalya,Kolosova; Vasiliy,Marchenko; Alexander,Ryzhikov | | | 2017-01-06 | 2017-02-17 | 2017-02-11 |
| EPI_ISL_247724 | EPI909449\|A/w | EPI909450\|A/wil | EPI909451\|A/wild | EPI909452\|A/wild | EPI909453\|A/wild | EPI909454\|A/wild | EPI909455\|A/wild | EPI909456\|A/wild | A/wild duck/Tatarstan/3059/2016 | A / H5N8 | e2 | Europe / Russ | Duck | Ivan Susloparov (State Research Center of Virology and Biotechnology (VECTOR) | State Research Center of Virology and Biotechnology (VECTOR | | Ivan,Susloparov; Natalya,Goncharova; Natalya,Kolosova; Vasiliy,Marchenko; Alexander,Ryzhikov | | | 2016-10-02 | 2017-02-12 | 2017-02-12 |
| EPI_ISL_247725 | EPI909457\|A/c | EPI909458\|A/chi | EPI909459\|A/chicl | EPI909460\|A/chicl | EPI909461\|A/chicl | EPI909462\|A/chicl | EPI909463\|A/chicl | EPI909464\|A/chicl | A/chicken/Kalmykia/2643/2016 | A / H5N8 | e2 | Europe / Russ | Gallus gallus | Ivan Susloparov (State Research Center of Virology and Biotechnology (VECTOR) | State Research Center of Virology and Biotechnology (VECTOR | | Ivan,Susloparov; Natalya,Goncharova; Natalya,Kolosova; Vasiliy,Marchenko; Alexander,Ryzhikov | | | 2016-11-21 | 2017-02-12 | 2017-02-12 |
| EPI_ISL_255181 | EPI954556\|A/s | EPI954557\|A/sw | EPI954555\|A/swa | EPI954559\|A/swa | EPI954552\|A/swa | EPI954558\|A/swa | EPI954554\|A/swa | EPI954553\|A/swa | A/swan/Italy/17VIR537-2/2017 | A / H5N8 | | Europe / Italy | Swan | Bianca Zecchin (Istituto Zooprofilattico Sperimentale Delle Venezie) | Istituto Zooprofilattico Sperimentale Delle Venezie | 17VIR537-2 | Zecchin, B.; Fusaro, A.; Milani, A.; Schivo, A.; Salviato, A.; Za | Istituto Zooprofilattico Sperimentale Delle Venezie | 17VIR537-2 | 2017-01-19 | 2017-03-13 | 2017-03-13 |
| EPI_ISL_255182 | EPI954564\|A/t | EPI954565\|A/tur | EPI954563\|A/turke | EPI954567\|A/turke | EPI954560\|A/turke | EPI954566\|A/turke | EPI954562\|A/turke | EPI954561\|A/turke | A/turkey/Italy/17VIR538-1/2017 | A / H5N8 | | Europe / Italy | Turkey | Bianca Zecchin (Istituto Zooprofilattico Sperimentale Delle Venezie) | Istituto Zooprofilattico Sperimentale Delle Venezie | 17VIR538-1 | Zecchin, B.; Fusaro, A.; Milani, A.; Schivo, A.; Salviato, A.; Za | Istituto Zooprofilattico Sperimentale Delle Venezie | 17VIR538-1 | 2017-01-20 | 2017-03-13 | 2017-03-13 |
| EPI_ISL_255194 | EPI954660\|A/( | EPI954661\|A/Go | EPI954659\|A/Goo | EPI954663\|A/Goo | EPI954656\|A/Goo | EPI954662\|A/Goo | EPI954658\|A/Goo | EPI954657\|A/Goo | A/Goose/Hungary/1030/2017 | A / H5N8 | c1 | Europe / Hung | Avian | Adam Dan (Danam.Vet.Molbiol) | Danam.Vet.Molbiol | 1030 | Adam, Dan | Danam.Vet.Molbiol | 1030 | 2017-01-11 | 2017-03-22 | 2017-03-13 |
| EPI_ISL_255206 | EPI954756\|A/I | EPI954757\|A/Pe | EPI954755\|A/Pere | EPI954759\|A/Pere | EPI954752\|A/Pere | EPI954758\|A/Pere | EPI954754\|A/Pere | EPI954753\|A/Pere | A/Peregrine_falcon/Hungary/4882/2017 | A / H5N8 | c1 | Europe / Hung | Falco peregrir | Adam Dan (Danam.Vet.Molbiol) | Danam.Vet.Molbiol | 4882 | Adam, Dan | Danam.Vet.Molbiol | 4882 | 2017-02-07 | 2017-03-14 | 2017-03-13 |
| EPI_ISL_255212 | EPI954810\|A/( | EPI954811\|A/Gr | EPI954809\|A/Grey | EPI954813\|A/Grey | EPI954806\|A/Grey | EPI954812\|A/Grey | EPI954808\|A/Grey | EPI954807\|A/Grey | A/Greylag_goose/Hungary/320/2017 | A / H5N8 | c1 | Europe / Hung | Anser anser | Adam Dan (Danam.Vet.Molbiol) | Danam.Vet.Molbiol | 320 | Adam, Dan | Danam.Vet.Molbiol | 320 | 2017-01-04 | 2017-03-22 | 2017-03-13 |
| EPI_ISL_255220 | EPI954874\|A/I | EPI954875\|A/Ma | EPI954873\|A/Malla | EPI954877\|A/Malla | EPI954870\|A/Malla | EPI954876\|A/Malla | EPI954872\|A/Malla | EPI954871\|A/Malla | A/Mallard/Hungary/1574a/2017 | A / H5N8 | c1 | Europe / Hung | Anas platyrhy | Adam Dan (Danam.Vet.Molbiol) | Danam.Vet.Molbiol | 1574a | Adam, Dan | Danam.Vet.Molbiol | 1574a | 2017-01-14 | 2017-03-22 | 2017-03-13 |
| EPI_ISL_266536 | EPI1007664\|A | EPI1007665\|A/cl | EPI1007666\|A/chi | EPI1007667\|A/chi | EPI1007668\|A/chi | EPI1007669\|A/chi | EPI1007670\|A/chi | EPI1007671\|A/chi | A/chicken/Belgium/807/2017 | A / H5N8 | passage detai | Europe / Belgi | Gallus gallus | | | | Steensels,M.; Lambrecht,B.; Vandenbussche,F.; Van Borm,S. | | | 2017-02-01 | | 2017-05-10 |
| EPI_ISL_266537 | EPI1007672\|A | EPI1007673\|A/p | EPI1007674\|A/pea | EPI1007675\|A/pea | EPI1007676\|A/pea | EPI1007677\|A/pea | EPI1007678\|A/pea | EPI1007679\|A/pea | A/peacock/Belgium/1017/2017 | A / H5N8 | passage detai | Europe / Belgi | Pavo cristatus | | | | Steensels,M.; Lambrecht,B.; Vandenbussche,F.; Van Borm,S. | | | 2017-02-08 | | 2017-05-10 |
| EPI_ISL_266820 | EPI1009533\|A | EPI1009534\|A/B | EPI1009535\|A/Bea | EPI1009536\|A/Bea | EPI1009537\|A/Bea | EPI1009538\|A/Bea | EPI1209290\|A/Bea | EPI1009540\|A/Bea | A/Bean goose/Hubei/CH-i122/2017_H5N8 | A / H5N8 | egg2 | Asia / China / | Anser fabalis | Jianjun Chen (Wuhan Institute of Virology / Chinese Academy of Sciences) | Wuhan Institute of Virology | | | | | 2017-02-24 | 2017-07-22 | 2017-06-13 |
| EPI_ISL_268513 | EPI1018204\|A | EPI1018205\|A/d | EPI1018206\|A/duc | EPI1018207\|A/duc | EPI1018208\|A/duc | EPI1018209\|A/duc | EPI1018210\|A/duc | EPI1018211\|A/duc | A/duck/Egypt/F446/2017 | A / H5N8 | E1 | Africa / Egypt | Duck | Abdel-Satar Mohamed Arafa (Animal Health Research Institute) | Animal Health Research Institute | F446 | 'Nahed,Yehia; Mahmoud,Naguib; Naglaa,Hagag; Mohamed,E | National Laboratory for Veterinary Quality Control on Po | | 2017-04-06 | 2017-06-29 | 2017-06-29 |
| EPI_ISL_268622 | EPI1019395\|A | EPI1019396\|A/cl | EPI1019394\|A/chi | EPI1019398\|A/chi | EPI1019391\|A/chi | EPI1019397\|A/chi | EPI1019393\|A/chi | EPI1019392\|A/chi | A/Ch/NL-Abbega/X16015736/2016 | A / H5N8 | Original | Europe / Neth | Gallus gallus | Saskia Bergervoet (Wageningen Bioveterinary Research) | Wageningen Bioveterinary Research | | Beerens, Nancy; Heutink, Rene; Harders, Frank; Verschuren-F | Wageningen Bioveterinary Research | X16015736 | 2016-12-12 | 2017-07-04 | 2017-07-04 |
| EPI_ISL_268623 | EPI1019403\|A | EPI1019404\|A/cl | EPI1019402\|A/chi | EPI1019406\|A/chi | EPI1019399\|A/chi | EPI1019405\|A/chi | EPI1019401\|A/chi | EPI1019400\|A/chi | A/Ch/NL-Boven Leeuwen/16016151-006-010/2016 | A / H5N8 | Original | Europe / Neth | Gallus gallus | Saskia Bergervoet (Wageningen Bioveterinary Research) | Wageningen Bioveterinary Research | | Beerens, Nancy; Heutink, Rene; Harders, Frank; Verschuren-F | Wageningen Bioveterinary Research | 16016151-00( | 2016-12-19 | 2017-07-04 | 2017-07-04 |
| EPI_ISL_268636 | EPI1019507\|A | EPI1019508\|A/e | EPI1019506\|A/eur | EPI1019510\|A/eur | EPI1019503\|A/eur | EPI1019509\|A/eur | EPI1019505\|A/eur | EPI1019504\|A/eur | A/Eur_Wig/NL-De Waal (Texel)/16014891-003/2016 | A / H5N8 | Original | Europe / Neth | Anas penelope | Saskia Bergervoet (Wageningen Bioveterinary Research) | Wageningen Bioveterinary Research | | Beerens, Nancy; Heutink, Rene; Harders, Frank; Verschuren-F | Wageningen Bioveterinary Research | 16014891-00: | 2016-11-27 | 2017-07-04 | 2017-07-04 |
| EPI_ISL_268638 | EPI1019523\|A | EPI1019524\|A/e | EPI1019522\|A/eur | EPI1019526\|A/eur | EPI1019519\|A/eur | EPI1019525\|A/eur | EPI1019521\|A/eur | EPI1019520\|A/eur | A/Eur_Wig/NL-Drieborg (Dollard)/16015513-001/2016 | A / H5N8 | Original | Europe / Neth | Anas penelope | Saskia Bergervoet (Wageningen Bioveterinary Research) | Wageningen Bioveterinary Research | | Beerens, Nancy; Heutink, Rene; Harders, Frank; Verschuren-F | Wageningen Bioveterinary Research | 16015513-00 | 2016-12-06 | 2017-07-04 | 2017-07-04 |
| EPI_ISL_268642 | EPI1019555\|A | EPI1019556\|A/e | EPI1019554\|A/eur | EPI1019558\|A/eur | EPI1019551\|A/eur | EPI1019557\|A/eur | EPI1019553\|A/eur | EPI1019552\|A/eur | A/Eur_Wig/NL-Greonterp/16015653-001/2016 | A / H5N8 | Original | Europe / Neth | Anas penelope | Saskia Bergervoet (Wageningen Bioveterinary Research) | Wageningen Bioveterinary Research | | Beerens, Nancy; Heutink, Rene; Harders, Frank; Verschuren-F | Wageningen Bioveterinary Research | 16015653-00 | 2016-12-08 | 2017-07-04 | 2017-07-04 |
| EPI_ISL_268652 | EPI1019635\|A | EPI1019636\|A/e | EPI1019634\|A/eur | EPI1019638\|A/eur | EPI1019631\|A/eur | EPI1019637\|A/eur | EPI1019633\|A/eur | EPI1019632\|A/eur | A/Eur_Wig/NL-Zoeterwoude/16015702-010/2016 | A / H5N8 | Original | Europe / Neth | Anas penelope | Saskia Bergervoet (Wageningen Bioveterinary Research) | Wageningen Bioveterinary Research | | Beerens, Nancy; Heutink, Rene; Harders, Frank; Verschuren-F | Wageningen Bioveterinary Research | 16015702-01( | 2016-12-10 | 2017-07-04 | 2017-07-04 |
| EPI_ISL_268679 | EPI1019851\|A | EPI1019852\|A/tu | EPI1019850\|A/tuft | EPI1019854\|A/tuft | EPI1019847\|A/tuft | EPI1019853\|A/tuft | EPI1019849\|A/tuft | EPI1019848\|A/tuft | A/T_Dk/NL-Zeewolde/16013976-006/2016 | A / H5N8 | Original | Europe / Neth | Aythya fuligula | Saskia Bergervoet (Wageningen Bioveterinary Research) | Wageningen Bioveterinary Research | | Beerens, Nancy; Heutink, Rene; Harders, Frank; Verschuren-F | Wageningen Bioveterinary Research | 16013976-00( | 2016-11-09 | 2017-07-04 | 2017-07-04 |
| EPI_ISL_268968 | | | | EPI1021124\|A/chi | | | | | A/chicken/Czech Republic/1689-17/2017 (H5N8) | A / H5N8 | | Europe / Czec | Gallus gallus | Alexander Nagy (State Veterinary Institute Prague) | State Veterinary Institute Prague | | Nagy,A | State Veterinary Institute Prague | | 2017-01-04 | 2017-07-06 | 2017-07-06 |
| EPI_ISL_271714 | EPI1032551\|A | EPI1032550\|A/M | EPI1032555\|A/Mu | EPI1032549\|A/Mu | EPI1032552\|A/Mu | EPI1032554\|A/Mu | EPI1032553\|A/Mu | EPI1032556\|A/Mu | A/Mulard_duck/Hungary/59163/2016 | A / H5N8 | tissues | Europe / Hung | Anas sp. | Adam Dan (Danam.Vet.Molbiol) | Danam.Vet.Molbiol | | Adam, Dan | National Food Chain Safety Office Veterinary Diagnosti | 59163 | 2016-11-24 | 2018-02-01 | 2017-07-26 |
| EPI_ISL_275288 | EPI1045614\|A | EPI1045615\|A/cl | EPI1045616\|A/chi | EPI1045619\|A/chi | EPI1045617\|A/chi | EPI1045618\|A/chi | EPI1045620\|A/chi | EPI1045621\|A/chi | A/chicken/Tatarstan/88/2017 | A / H5N8 | E2 | Europe / Russ | Gallus gallus | Ivan Susloparov (State Research Center of Virology and Biotechnology (VECTOR) | State Research Center of Virology and Biotechnology (VECTOR | | Ivan,Susloparov; Natalya,Goncharova; Natalya,Kolosova; Vasiliy,Marchenko; Alexander,Ryzhikov | | | 2017-05-08 | 2017-08-22 | 2017-08-22 |
| EPI_ISL_278200 | | | | EPI1061370\|A/Bra | | | | | A/Brahma chicken/Belgium/6153/2017 | A / H5N8 | passage detai | Europe / Belgi | Gallus gallus | | | | Steensels,M.; Lambrecht,B.; Vandenbussche,F.; Van Borm,S. | | | 2017-06-15 | | 2017-07-03 |
| EPI_ISL_284004 | EPI1187455\|A | EPI1106512\|A/C | EPI1106479\|A/Chi | EPI1093378\|A/Chi | EPI1187478\|A/Chi | EPI1093390\|A/Chi | EPI1187527\|A/Chi | EPI1187528\|A/Chi | A/Chicken/South_Africa/S2017/08_0336_P3/2017 | A / H5N8 | | Africa / South | Gallus gallus c | Florette Kathleen Treurnicht (National Institute for Communicable Diseases / Center | National Institute for Communicable Diseases | A/Chicken/Sou | | Western Cape Provincial Veterinary Laboratory | A/Chicken/Sou | 2017-08-21 | 2018-02-27 | 2017-11-13 |
| EPI_ISL_284683 | | | | EPI1098487\|A/Ost | EPI1187540\|A/Ost | EPI1098489\|A/Ost | EPI1187539\|A/Ost | EPI1187538\|A/Ost | A/Ostrich/South_Africa/S2017/08_0268_P2/2017 | A / H5N8 | | Africa / South | Ostrich | Florette Kathleen Treurnicht (National Institute for Communicable Diseases / Center | National Institute for Communicable Diseases | A/Ostrich/Sou | | Western Cape Provincial Veterinary Laboratory | A/Ostrich/Sou | 2017-08-16 | 2018-02-27 | 2017-11-20 |
| EPI_ISL_285512 | | | EPI1106462\|A/Chi | EPI1103669\|A/Chi | EPI1187740\|A/Chi | EPI1103675\|A/Chi | EPI1187732\|A/Chi | EPI1187730\|A/Chi | A/Chicken/South_Africa/S2017/08_0336_P1/2017 | A / H5N8 | | Africa / South | Gallus gallus c | Florette Kathleen Treurnicht (National Institute for Communicable Diseases / Center | National Institute for Communicable Diseases | A/Chicken/Sou | | Western Cape Provincial Veterinary Laboratory | A/Chicken/Sou | 2017-08-21 | 2018-01-10 | 2017-11-21 |
| EPI_ISL_287800 | EPI1119037\|A | EPI1119048\|A/s | EPI1119057\|A/spo | EPI1119065\|A/spo | EPI1119072\|A/spo | EPI1119073\|A/spo | EPI1119074\|A/spo | EPI1119075\|A/spo | A/spoonbill/Taiwan/DB645/2017 | A / H5N6 | organs | Asia / Taiwan | Other avian | Yu-Pin Liu (Animal Health Research Institute) | Animal Health Research Institute | | Yu-Ju,Lin; Li-Hsuan,Chen; Wan-Chen,Li; Yen-Ping,Chen; Yu-Pin,Liu; Fan,Lee; Wen-Jane,Tu | | | 2017-12-01 | 2017-12-12 | 2017-12-12 |
| EPI_ISL_287906 | EPI1119859\|A | EPI1119860\|A/D | EPI1119861\|A/Du | EPI1119862\|A/Du | EPI1119863\|A/Du | EPI1119864\|A/Du | EPI1119865\|A/Du | EPI1119866\|A/Du | A/Duck/Netherlands/17017236-001-005/2017 | A / H5N6 | Original | Europe / Neth | Duck | Saskia Bergervoet (Wageningen Bioveterinary Research) | Wageningen Bioveterinary Research | | Beerens, Nancy; Heutink, Rene; Harders, Frank; Verschuren-F | Wageningen Bioveterinary Research | 17017236-00 | 2017-12-07 | 2017-12-12 | 2017-12-12 |
| EPI_ISL_287907 | EPI1119867\|A | EPI1119868\|A/D | EPI1119869\|A/Du | EPI1119870\|A/Du | EPI1119871\|A/Du | EPI1119872\|A/Du | EPI1119873\|A/Du | EPI1119874\|A/Du | A/Duck/Netherlands/17017237-001-005/2017 | A / H5N6 | Original | Europe / Neth | Duck | Saskia Bergervoet (Wageningen Bioveterinary Research) | Wageningen Bioveterinary Research | | Beerens, Nancy; Heutink, Rene; Harders, Frank; Verschuren-F | Wageningen Bioveterinary Research | 17017237-00 | 2017-12-07 | 2017-12-12 | 2017-12-12 |
| EPI_ISL_288362 | EPI1122874\|A | EPI1122875\|A/cl | EPI1122876\|A/chi | EPI1122877\|A/chi | EPI1122878\|A/chi | EPI1122879\|A/chi | EPI1122880\|A/chi | EPI1122881\|A/chi | A/chicken/Greece/39_2017/2017 | A / H5N6 | Egg Passage | Europe / Gree | Gallus gallus c | James Seekings (Animal and Plant Health Agency (APHA) / Virology Department) | Animal and Plant Health Agency (APHA) | AV000624-17 | Seekings, James; Ellis, Richard; Brookes, Sharon M; Reid, St | Thessalonica Veterinary Centre (TVC) | 39/2017 | 2017-02-06 | 2017-12-14 | 2017-12-14 |
| EPI_ISL_288364 | EPI1122890\|A | EPI1122891\|A/cl | EPI1122892\|A/chi | EPI1122893\|A/chi | EPI1122894\|A/chi | EPI1122895\|A/chi | EPI1122896\|A/chi | EPI1122897\|A/chi | A/chicken/Greece/39_2017b/2017 | A / H5N6 | Egg Passage | Europe / Gree | Gallus gallus c | James Seekings (Animal and Plant Health Agency (APHA) / Virology Department) | Animal and Plant Health Agency (APHA) | AV000624-17 | Seekings, James; Ellis, Richard; Brookes, Sharon M; Essen, S | Thessalonica Veterinary Centre (TVC) | 39/2017b | 2017-02-06 | 2017-12-14 | 2017-12-14 |
| EPI_ISL_288409 | EPI1123234\|A | EPI1123235\|A/m | EPI1123233\|A/mu | EPI1123237\|A/mu | EPI1123230\|A/mu | EPI1123236\|A/mu | EPI1123232\|A/mu | EPI1123231\|A/mu | A/Mute Swan/Netherlands/17017367-012/2017 | A / H5N6 | Original | Europe / Neth | Cygnus olor | Saskia Bergervoet (Wageningen Bioveterinary Research) | Wageningen Bioveterinary Research | | Beerens, Nancy; Heutink, Rene; Harders, Frank; Verschuren-F | Wageningen Bioveterinary Research | 17017367-01: | 2017-12-09 | 2017-12-14 | 2017-12-14 |
| EPI_ISL_288410 | EPI1123242\|A | EPI1123243\|A/m | EPI1123241\|A/mu | EPI1123245\|A/mu | EPI1123238\|A/mu | EPI1123244\|A/mu | EPI1123240\|A/mu | EPI1123239\|A/mu | A/Mute Swan/Netherlands/17017377-001/2017 | A / H5N6 | Original | Europe / Neth | Cygnus olor | Saskia Bergervoet (Wageningen Bioveterinary Research) | Wageningen Bioveterinary Research | | Beerens, Nancy; Heutink, Rene; Harders, Frank; Verschuren-F | Wageningen Bioveterinary Research | 17017377-00 | 2017-12-09 | 2017-12-14 | 2017-12-14 |
| EPI_ISL_288412 | EPI1123258\|A | EPI1123259\|A/T | EPI1123257\|A/T_I | EPI1123261\|A/T_I | EPI1123254\|A/T_I | EPI1123260\|A/T_I | EPI1123256\|A/T_I | EPI1123255\|A/T_I | A/Tufted Duck/Netherlands/17017367-007/2017 | A / H5N6 | Original | Europe / Neth | Aythya fuligula | Saskia Bergervoet (Wageningen Bioveterinary Research) | Wageningen Bioveterinary Research | | Beerens, Nancy; Heutink, Rene; Harders, Frank; Verschuren-F | Wageningen Bioveterinary Research | 17017367-00 | 2017-12-09 | 2017-12-14 | 2017-12-14 |
| EPI_ISL_288438 | EPI1123340\|A | EPI1123341\|A/cl | EPI1123342\|A/chi | EPI1123343\|A/chi | EPI1123344\|A/chi | EPI1123345\|A/chi | EPI1123346\|A/chi | EPI1123347\|A/chi | A/chicken/Korea/Gunsan/2017 | A / H5N8 | c1 | Asia / Korea, | Gallus gallus c | Eun-Kyoung Lee (Animal and Plant Quarantine Agency (S-2026)) | Animal and Plant Quarantine Agency (S-2145) | | | Animal and Plant Quarantine Agency (O-2144) | | 2017-06-03 | 2017-12-17 | 2017-12-17 |
| EPI_ISL_289173 | EPI1127535\|A | EPI1127536\|A/m | EPI1127537\|A/mu | EPI1127538\|A/mu | EPI1127539\|A/mu | EPI1127540\|A/mu | EPI1127541\|A/mu | EPI1127556\|A/mu | A/mute swan/Shimane/3211A001/2017 | A / H5N6 | | Asia / Japan / | Cygnus olor | | | | Soda,K.; Ito,H.; Usui,T.; Ozaki,H.; Murase,T.; Yamaguchi,T.; Ito,T. | | | 2017-11-05 | | 2017-11-19 |
| EPI_ISL_292334 | EPI1145114\|A | EPI1145115\|A/B | EPI1145116\|A/Ba | EPI1145117\|A/Ba | EPI1145118\|A/Ba | EPI1145119\|A/Ba | EPI1145120\|A/Ba | EPI1145121\|A/Ba | A/Bar-headed Goose/Qinghai/B655/2017 | A / H5N8 | egg2 | Asia / China / | Anser indicus | Jianjun Chen (Wuhan Institute of Virology / Chinese Academy of Sciences) | Wuhan Institute of Virology | | | Wuhan Institute of Virology, CAS | | 2017-05-17 | 2018-01-22 | 2018-01-22 |
| EPI_ISL_297235 | EPI1169196\|A | EPI1169197\|A/cl | EPI1169198\|A/chi | EPI1169201\|A/chi | EPI1169199\|A/chi | EPI1169200\|A/chi | EPI1169202\|A/chi | EPI1169203\|A/chi | A/chicken/Rostov-on-Don/1598/2017 | A / H5N8 | Original | Europe / Russ | Gallus gallus c | Ivan Susloparov (State Research Center of Virology and Biotechnology (VECTOR) | State Research Center of Virology and Biotechnology (VECTOR | | Ivan,Susloparov; Natalya,Goncharova; Natalya,Kolosova; Vasiliy,Marchenko; Alexander,Ryzhikov | | | 2017-11-17 | 2018-02-09 | 2018-02-07 |
| EPI_ISL_298648 | | | | EPI1176525\|A/gre | | | | | A/great egret/Israel/1088/2016 | A / H5N8 | passage detai | Asia / Israel | Other avian | | | | Shkoda,I.; Lapin,K.; Simanov,L.; Lublin,A. | | | 2016-12-25 | | 2017-05-28 |
| EPI_ISL_298649 | | | | EPI1176526\|A/turk | | | | | A/turkey/Israel/184/2017 | A / H5N8 | passage detai | Asia / Israel | Turkey | | | | Shkoda,I.; Lapin,K.; Simanov,L.; Lublin,A. | | | 2017-02-05 | | 2017-05-28 |
| EPI_ISL_304404 | EPI1202731\|A | EPI1202730\|A/F | EPI1202724\|A/Fuj | EPI1202729\|A/Fuj | EPI1202726\|A/Fuj | EPI1202727\|A/Fuj | EPI1202728\|A/Fuj | EPI1202725\|A/Fuj | A/Fujian-Sanyuan/21099/2017 | A / H5N6 | C1 | Asia / China / | Human | Lei Yang (WHO Chinese National Influenza Center / Virology Institute, Chinese CDC | WHO Chinese National Influenza Center | | | Fujian Provincial Center for Disease Control and Prevent | | 2017-12-25 | 2018-03-20 | 2018-03-20 |
| EPI_ISL_308783 | EPI1223758\|A | EPI1223759\|A/ju | EPI1223760\|A/jung | EPI1223761\|A/jung | EPI1223762\|A/jung | EPI1223763\|A/jung | EPI1223789\|A/jung | EPI1223790\|A/jung | A/jungle crow/Hyogo/2803E022/2018 | A / H5N6 | | Asia / Japan / | Corvus macro | | | | Soda,K.; Ito,H.; Ozaki,H.; Usui,T.; Murase,T.; Yamaguchi,T.; Ito,T. | | | 2018-03-06 | | 2018-04-05 |
| EPI_ISL_309195 | EPI1226459\|A | EPI1226458\|A/C | EPI1226457\|A/Cyg | EPI1226452\|A/Cyg | EPI1226455\|A/Cyg | EPI1226454\|A/Cyg | EPI1226453\|A/Cyg | EPI1226456\|A/Cyg | A/Cygnus olor/Belgium/2967/2017 | A / H5N8 | passage detai | Europe / Belgi | Cygnus olor | | | | Van Borm,S.; Vandenbussche,F.; Mathijs,E.; Lambrecht,B.; Steensels,M. | | | 2017-03-21 | | 2018-03-23 |
| EPI_ISL_309197 | EPI1226476\|A | EPI1226475\|A/B | EPI1226474\|A/Bra | EPI1226468\|A/Bra | EPI1226472\|A/Bra | EPI1226471\|A/Bra | EPI1226470\|A/Bra | EPI1226473\|A/Bra | A/Brahma chicken/Belgium/6153/2017 | A / H5N8 | passage detai | Europe / Belgi | Chicken | | | | Van Borm,S.; Vandenbussche,F.; Mathijs,E.; Lambrecht,B.; Steensels,M. | | | 2017-06-15 | | 2018-03-23 |
| EPI_ISL_316531 | EPI1261492\|P | EPI1261493\|PB | EPI1261491\|PA_1 | EPI1261495\|HA_1 | EPI1261488\|NP_1 | EPI1261494\|NA_1 | EPI1261490\|MP_ | EPI1261489\|NS_1 | A/pheasant/Italy/17VIR7065-21/2017 | A / H5N8 | | Europe / Italy | Pheasant | Bianca Zecchin (Istituto Zooprofilattico Sperimentale Delle Venezie) | Istituto Zooprofilattico Sperimentale Delle Venezie | | Mulatti, P.; Fusaro, A.; Scolamacchia, F.; Zecchin, B.; Azzolini | Istituto Zooprofilattico Sperimentale Delle Venezie | | 2017-08-19 | 2018-07-26 | 2018-07-26 |
| EPI_ISL_316539 | EPI1261556\|P | EPI1261557\|PB | EPI1261555\|PA_1 | EPI1261559\|HA_1 | EPI1261552\|NP_1 | EPI1261558\|NA_1 | EPI1261554\|MP_ | EPI1261553\|NS_1 | A/turkey/Italy/17VIR8275-3/2017 | A / H5N8 | | Europe / Italy | Turkey | Bianca Zecchin (Istituto Zooprofilattico Sperimentale Delle Venezie) | Istituto Zooprofilattico Sperimentale Delle Venezie | | Mulatti, P.; Fusaro, A.; Scolamacchia, F.; Zecchin, B.; Azzolini | Istituto Zooprofilattico Sperimentale Delle Venezie | | 2017-09-26 | 2018-07-26 | 2018-07-26 |
| EPI_ISL_316553 | EPI1261667\|P | EPI1261668\|PB | EPI1261666\|PA_1 | EPI1261670\|HA_1 | EPI1261663\|NP_1 | EPI1261669\|NA_1 | EPI1261665\|MP_ | EPI1261664\|NS_1 | A/chicken/Italy/17VIR9113/2017 | A / H5N8 | | Europe / Italy | Chicken | Bianca Zecchin (Istituto Zooprofilattico Sperimentale Delle Venezie) | Istituto Zooprofilattico Sperimentale Delle Venezie | | Mulatti, P.; Fusaro, A.; Scolamacchia, F.; Zecchin, B.; Azzolini | Istituto Zooprofilattico Sperimentale Delle Venezie | | 2017-10-16 | 2018-07-26 | 2018-07-26 |
| EPI_ISL_316581 | EPI1261890\|P | EPI1261891\|PB | EPI1261889\|PA_1 | EPI1261893\|HA_1 | EPI1261886\|NP_1 | EPI1261892\|NA_1 | EPI1261888\|MP_ | EPI1261887\|NS_1 | A/duck/Italy/17VIR10767-13/2017 | A / H5N8 | | Europe / Italy | Duck | Bianca Zecchin (Istituto Zooprofilattico Sperimentale Delle Venezie) | Istituto Zooprofilattico Sperimentale Delle Venezie | | Mulatti, P.; Fusaro, A.; Scolamacchia, F.; Zecchin, B.; Azzolini | Istituto Zooprofilattico Sperimentale Delle Venezie | | 2017-12-01 | 2018-07-26 | 2018-07-26 |
| EPI_ISL_316582 | EPI1261898\|P | EPI1261899\|PB | EPI1261897\|PA_1 | EPI1261901\|HA_1 | EPI1261894\|NP_1 | EPI1261900\|NA_1 | EPI1261896\|MP_ | EPI1261895\|NS_1 | A/turkey/Italy/17VIR10983-22/2017 | A / H5N8 | | Europe / Italy | Turkey | Bianca Zecchin (Istituto Zooprofilattico Sperimentale Delle Venezie) | Istituto Zooprofilattico Sperimentale Delle Venezie | | Mulatti, P.; Fusaro, A.; Scolamacchia, F.; Zecchin, B.; Azzolini | Istituto Zooprofilattico Sperimentale Delle Venezie | | 2017-12-11 | 2018-07-26 | 2018-07-26 |
| EPI_ISL_320607 | EPI1270726\|A | EPI1270727\|A/cl | EPI1270728\|A/chi | EPI1270724\|A/chi | EPI1270729\|A/chi | EPI1270725\|A/chi | EPI1270730\|A/chi | EPI1270731\|A/chi | A/chicken/Kursk/284/2018 | A / H5N8 | E2 | Europe / Russ | Gallus gallus c | Ivan Susloparov (State Research Center of Virology and Biotechnology (VECTOR) | State Research Center of Virology and Biotechnology (VECTOR | | Alexey,Danilenko; Vasiliy,Marchenko; Ivan,Susloparov; Natalya,Goncharova; Natalya,Kolosova; Juliya, Bulanovich; Ale | | | 2018-06-11 | 2018-08-20 | 2018-08-20 |
| EPI_ISL_322179 | EPI1279362\|A | EPI1279363\|A/d | EPI1279364\|A/dor | EPI1279361\|A/dor | EPI1279365\|A/dor | EPI1279366\|A/dor | EPI1279367\|A/dor | EPI1279368\|A/dor | A/domestic duck/Germany-MV/AR613-L02727/2018 | A / H5N6 | | Europe / Gern | Anas platyrhy | Jacqueline King (Friedrich-Loeffler-Institut) | Friedrich-Loeffler-Institut | AR613-18 | | | | 2018-08-31 | 2018-09-07 | 2018-09-07 |
| EPI_ISL_332430 | EPI1327053\|A | EPI1327054\|A/cl | EPI1327052\|A/chi | EPI1327056\|A/chi | EPI1327049\|A/chi | EPI1327055\|A/chi | EPI1327051\|A/chi | EPI1327050\|A/chi | A/Chicken/Netherlands/18003041-001-005/2018 | A / H5N6 | Original | Europe / Neth | Gallus gallus | Saskia Bergervoet (Wageningen Bioveterinary Research) | Wageningen Bioveterinary Research | 18003041-00 | Beerens, Nancy; Heutink, Rene; Harders, Frank; Verschuren-F | Wageningen Bioveterinary Research | | 2018-02-25 | 2018-12-11 | 2018-12-11 |
| EPI_ISL_332804 | EPI1328853\|3( | EPI1328854\|300 | EPI1328852\|3000( | EPI1328856\|3000( | EPI1328849\|3000( | EPI1328855\|3000( | EPI1328851\|3000( | EPI1328850\|3000( | A/duck/Bangladesh/19D857/2017 | A / H5N6 | Original | Asia / Banglac | Avian | Yunho Jang (Centers for Disease Control and Prevention / WHO Collaborating Cent | Centers for Disease Control and Prevention | 3000682224 | Yang, G.; Chowdury, S; Hodges, E; Rahman, M.Z.; Jang, Y.; I | icddr,b International Centre for Diarrhoeal Disease Rese | 19D857 | 2017-05-03 | 2018-12-18 | 2018-12-18 |
| EPI_ISL_333133 | EPI1330540\|3( | EPI1330541\|300 | EPI1330539\|3000( | EPI1330543\|3000( | EPI1330536\|3000( | EPI1330542\|3000( | EPI1330538\|3000( | EPI1330537\|3000( | A/duck/Bangladesh/19D770/2017 | A / H5N6 | Original | Asia / Banglac | Avian | Yunho Jang (Centers for Disease Control and Prevention / WHO Collaborating Cent | Centers for Disease Control and Prevention | 3000682193 | Yang, G.; Chowdury, S; Hodges, E; Rahman, M.Z.; Jang, Y.; I | icddr,b International Centre for Diarrhoeal Disease Rese | 19D770 | 2017-01-11 | 2018-12-19 | 2018-12-19 |
| EPI_ISL_333629 | EPI1333866\|A | EPI1333867\|A/E | EPI1333868\|A/Em | EPI1333871\|A/Em | EPI1333870\|A/Em | EPI1333872\|A/Em | EPI1333873\|A/Em | EPI1333883\|A/Em | A/Environment/Fujiansanyuan/08/2017 | A / H5N6 | | Asia / China / | Feces | Xie Jianfeng (Fujian Center for Disease Control and Prevention) | Fujian Center for Disease Control and Prevention | | | Fujian Provincial Center for Disease Control and Prevent | | 2017-12-26 | 2018-12-29 | 2018-12-29 |
| EPI_ISL_335454 | EPI1342076\|A | EPI1342077\|A/cl | EPI1342075\|A/chi | EPI1342079\|A/chi | EPI1342072\|A/chi | EPI1342078\|A/chi | EPI1342074\|A/chi | EPI1342073\|A/chi | A/chicken/Korea/H239/2018 | A / H5N6 | | Asia / Korea, | Chicken | Yu-Na Lee (Animal and Plant Quarantine Agency (S-2158) / Avian Influenza Researc | Animal and Plant Quarantine Agency (S-2158) | | | | | 2018-03-16 | 2019-01-17 | 2019-01-17 |
| EPI_ISL_336933 | EPI1351440\|A | EPI1351441\|A/V | EPI1351442\|A/Vo | EPI1351443\|A/Vo | EPI1351444\|A/Vo | EPI1351445\|A/Vo | EPI1351446\|A/Vo | EPI1351447\|A/Vo | A/chicken/Voronezh/1513/2018 | A / H5N8 | E1 | Europe / Russ | Gallus gallus c | Alexey Danilenko (State Research Center of Virology and Biotechnology (VECTOR) | State Research Center of Virology and Biotechnology (VECTOR | | Natalya,Goncharova; Alexey,Danilenko; Vasiliy,Marchenko; Ivan,Susloparov; Natalya,Kolosova; Juliya, Bulanovich; Ale | | | 2018-11-02 | 2019-02-06 | 2019-01-28 |
| EPI_ISL_337151 | EPI1352496\|A | EPI1352497\|A/cl | EPI1352498\|A/chi | EPI1352499\|A/chi | EPI1352500\|A/chi | EPI1352501\|A/chi | EPI1352502\|A/chi | EPI1352503\|A/chi | A/chicken/Mari El/870/2018 | A / H5N8 | E2 | Europe / Russ | Gallus gallus c | Natalia Goncharova (State Research Center of Virology and Biotechnology (VECTO | State Research Center of Virology and Biotechnology (VECTOR | | Natalia,Goncharova; Ivan,Susloparov; Natalia,Kolosova; Alexey,Danilenko; Juliya,Bulanovich; Vasiliy,Marchenko; Alexa | | | 2018-08-03 | 2019-02-18 | 2019-01-29 |
| EPI_ISL_344527 | EPI1420351\|A | EPI1420350\|A/D | EPI1420349\|A/Du | EPI1381391\|A/Du | EPI1420346\|A/Du | EPI1420345\|A/Du | EPI1420344\|A/Du | EPI1420347\|A/Du | A/Duck/Egypt/AR518/2017 | A / H5N8 | clinical | Africa / Egypt | Duck | Timm Harder (Friedrich-Loeffler-Institut) | Friedrich-Loeffler-Institut | | Hassan ,KE; King,J ; El-Kady, MF; Abohamra ,S; Pohlmann, | Poultry Diseases Department ,Faculty of Veterinary Med | | 2017-03-08 | 2019-04-15 | 2019-03-08 |
| EPI_ISL_344528 | | | | EPI1381392\|A/Chi | | | | | A/Chicken/Egypt/AR520/2017 | A / H5N8 | clinical | Africa / Egypt | Chicken | Timm Harder (Friedrich-Loeffler-Institut) | Friedrich-Loeffler-Institut | | Hassan KE, El-Khady MF, Abohamra S, Harder TC | | | 2017-03-10 | 2019-03-08 | 2019-03-08 |
| EPI_ISL_344531 | | | | EPI1381424\|A/Chi | | | | | A/Chicken/Egypt/AR539/2018 | A / H5N8 | clinical | Africa / Egypt | Chicken | Timm Harder (Friedrich-Loeffler-Institut) | Friedrich-Loeffler-Institut | | Hassan KE, El-Khady MF, Abohamra S, Harder TC | | | 2018-01-02 | 2019-03-08 | 2019-03-08 |
| EPI_ISL_344537 | | | | EPI1381427\|A/Chi | | | | | A/Chicken/Egypt/AR548/2018 | A / H5N8 | clinical | Africa / Egypt | Chicken | Timm Harder (Friedrich-Loeffler-Institut) | Friedrich-Loeffler-Institut | | Hassan KE, El-Khady MF, Abohamra S, Harder TC | | | 2018-03-20 | 2019-03-08 | 2019-03-08 |
| EPI_ISL_344538 | | | | EPI1381394\|A/Chi | | | | | A/Chicken/Egypt/AR549/2018 | A / H5N8 | clinical | Africa / Egypt | Chicken | Timm Harder (Friedrich-Loeffler-Institut) | Friedrich-Loeffler-Institut | | Hassan KE, El-Khady MF, Abohamra S, Harder TC | | | 2018-03-20 | 2019-03-08 | 2019-03-08 |
| EPI_ISL_344539 | EPI1420343\|A | EPI1420342\|A/T | EPI1420341\|A/Tur | EPI1381395\|A/Tur | EPI1420339\|A/Tur | EPI1420338\|A/Tur | EPI1420336\|A/Tur | EPI1420340\|A/Tur | A/Turkey/Egypt/AR550/2018 | A / H5N8 | clinical | Africa / Egypt | Turkey | Timm Harder (Friedrich-Loeffler-Institut) | Friedrich-Loeffler-Institut | | Hassan ,KE; King,J ; El-Kady, MF; Abohamra ,S; Pohlmann, | Poultry Diseases Department ,Faculty of Veterinary Med | | 2018-03-28 | 2019-10-11 | 2019-03-08 |
| EPI_ISL_344543 | | | | EPI1381399\|A/Chi | | | | | A/Chicken/Egypt/AR557/2018 | A / H5N8 | clinical | Africa / Egypt | Chicken | Timm Harder (Friedrich-Loeffler-Institut) | Friedrich-Loeffler-Institut | | Hassan KE, El-Khady MF, Abohamra S, Harder TC | | | 2018-04-25 | 2019-03-08 | 2019-03-08 |
| EPI_ISL_344544 | | | | EPI1381400\|A/Chi | | | | | A/Chicken/Egypt/AR558/2018 | A / H5N8 | clinical | Africa / Egypt | Chicken | Timm Harder (Friedrich-Loeffler-Institut) | Friedrich-Loeffler-Institut | | Hassan KE, El-Khady MF, Abohamra S, Harder TC | | | 2018-05-18 | 2019-03-08 | 2019-03-08 |
| EPI_ISL_344545 | | | | EPI1381401\|A/Chi | | | | | A/Chicken/Egypt/AR559/2018 | A / H5N8 | clinical | Africa / Egypt | Chicken | Timm Harder (Friedrich-Loeffler-Institut) | Friedrich-Loeffler-Institut | | Hassan KE, El-Khady MF, Abohamra S, Harder TC | | | 2018-05-20 | 2019-03-08 | 2019-03-08 |
| EPI_ISL_344546 | EPI1420334\|A | EPI1420333\|A/D | EPI1420332\|A/Du | EPI1381402\|A/Du | EPI1420330\|A/Du | EPI1420329\|A/Du | EPI1420328\|A/Du | EPI1420331\|A/Du | A/Duck/Egypt/AR560/2018 | A / H5N8 | clinical | Africa / Egypt | Duck | Timm Harder (Friedrich-Loeffler-Institut) | Friedrich-Loeffler-Institut | | Hassan ,KE; King,J ; El-Kady, MF; Abohamra ,S; Pohlmann, | Poultry Diseases Department ,Faculty of Veterinary Med | | 2018-05-25 | 2019-04-15 | 2019-03-08 |
| EPI_ISL_344547 | | | | EPI1381403\|A/Chi | | | | | A/Chicken/Egypt/AR562/2018 | A / H5N8 | 2nd passage | Africa / Egypt | Chicken | Timm Harder (Friedrich-Loeffler-Institut) | Friedrich-Loeffler-Institut | | Hassan KE, El-Khady MF, Abohamra S, Harder TC | | | 2018-06-25 | 2019-03-08 | 2019-03-08 |
| EPI_ISL_344548 | | | | EPI1381404\|A/Chi | | | | | A/Chicken/Egypt/AR591/2018 | A / H5N8 | clinical | Africa / Egypt | Chicken | Timm Harder (Friedrich-Loeffler-Institut) | Friedrich-Loeffler-Institut | | Hassan KE, El-Khady MF, Abohamra S, Harder TC | | | 2018-05-14 | 2019-03-08 | 2019-03-08 |
| EPI_ISL_366612 | | | | EPI1500993\|A/Po | | | | | A/Poultry/Iran/clade 2344/2018 | A / H5N8 | | Asia / Iran, Isl | Poultry | | | | Abdoli,A. | | | 2018-01-22 | | 2019-05-14 |
| EPI_ISL_369336 | EPI1514083\|A | EPI1514084\|A/cl | EPI1514085\|A/chi | EPI1514086\|A/chi | EPI1514087\|A/chi | EPI1514088\|A/chi | EPI1514089\|A/chi | EPI1514090\|A/chi | A/chicken/South Africa/Villiers/2017 | A / H5N8 | | Africa / South | Chicken | | | | Abolnik,C.; Rauff,D.L.; Pieterse,R.; Peyrot,B.; Choma,P.; Ebersohn,K.; Phiri,T. | | | 2017-06-19 | | 2018-04-04 |
| EPI_ISL_369337 | EPI1514091\|A | EPI1514092\|A/cl | EPI1514093\|A/chi | EPI1514094\|A/chi | EPI1514095\|A/chi | EPI1514096\|A/chi | EPI1514097\|A/chi | EPI1514098\|A/chi | A/chicken/South Africa/Standerton/2017 | A / H5N8 | | Africa / South | Chicken | | | | Abolnik,C.; Rauff,D.L.; Pieterse,R.; Peyrot,B.; Choma,P.; Ebersohn,K.; Phiri,T. | | | 2017-06-20 | | 2018-04-04 |
| EPI_ISL_369366 | EPI1514323\|A | EPI1514324\|A/o | EPI1514325\|A/ost | EPI1514326\|A/ost | EPI1514327\|A/ost | EPI1514328\|A/ost | EPI1514329\|A/ost | EPI1514330\|A/ost | A/ostrich/South Africa/002/2017 | A / H5N8 | | Africa / South | Ostrich | | | | Abolnik,C.; Rauff,D.L.; Pieterse,R.; Peyrot,B.; Choma,P.; Ebersohn,K.; Phiri,T. | | | 2017-10-26 | | 2018-04-04 |
| EPI_ISL_378362 | EPI1549838\|A | EPI1549839\|A/w | EPI1549840\|A/wh | EPI1549841\|A/wh | EPI1549842\|A/wh | EPI1549843\|A/wh | EPI1549844\|A/wh | EPI1549845\|A/wh | A/whooper swan/Shanxi/RC01/2016 | A / H5N8 | | Asia / China | Whooper swa | | | | Li,M.; Zhao,N.; Ma,J.; Feng,S.; Chang,H.; Luo,J.; He,H. | | | 2016-12-20 | | 2016-12-26 |
| EPI_ISL_378362 | EPI1549838\|A | EPI1549839\|A/w | EPI1549840\|A/wh | EPI1549841\|A/wh | EPI1549842\|A/wh | EPI1549843\|A/wh | EPI1549844\|A/wh | EPI1549845\|A/wh | A/whooper swan/Shanxi/RC01/2016 | A / H5N8 | | Asia / China | Whooper swa | | | | Li,M.; Zhao,N.; Ma,J.; Feng,S.; Chang,H.; Luo,J.; He,H. | | | 2016-12-20 | | 2016-12-26 |
| EPI_ISL_379982 | EPI1557366\|A | EPI1557369\|A/W | EPI1557360\|A/Wh | EPI1557352\|A/Wh | EPI1557354\|A/Wh | EPI1557372\|A/Wh | EPI1557363\|A/Wh | EPI1557357\|A/Wh | A/White-fronted Goose/AN/1-15-12/2016 | A / H5N8 | | Europe / Ukra | White-fronted | | | | Muzyka,D.; Rula,O.; Tkachenko,S.; Muzyka,N.; Kothe,S.; Stegniy,B.; Pantin-Jackwood,M.; Beer,M. | | | 2016-12-15 | | 2018-09-27 |
| EPI_ISL_379983 | EPI1557367\|A | EPI1557370\|A/R | EPI1557361\|A/Ru | EPI1557353\|A/Ru | EPI1557355\|A/Ru | EPI1557373\|A/Ru | EPI1557364\|A/Ru | EPI1557358\|A/Ru | A/Ruddy Shelduck/AN/2-14-12/2016 | A / H5N8 | | Europe / Ukra | Ruddy shelduc | | | | Muzyka,D.; Rula,O.; Tkachenko,S.; Muzyka,N.; Kothe,S.; Stegniy,B.; Pantin-Jackwood,M.; Beer,M. | | | 2016-12-14 | | 2018-09-27 |
| EPI_ISL_380991 | EPI1564190\|A | EPI1564191\|A/C | EPI1564192\|A/Cro | EPI1564193\|A/Cro | EPI1564194\|A/Cro | EPI1564195\|A/Cro | EPI1564196\|A/Cro | EPI1564197\|A/Cro | A/Crow/Aghakhan/2017 | A / H5N8 | | Asia / Iran, Isl | Crow | | | | Ghalyanchilangaroudi,A.; Ziafati Kafi,Z.; Malekan,M.; Fallah,M.H.; Ghafoori,A. | | | 2017-09-08 | | 2018-11-15 |
| EPI_ISL_381810 | EPI1567248\|A | EPI1567249\|A/E | EPI1567250\|A/Egy | EPI1567251\|A/Egy | | EPI1567252\|A/Egy | EPI1567253\|A/Egy | EPI1567254\|A/Egy | A/Egyptian goose/South Africa/001/2017 | A / H5N8 | | Africa / South | Egyptian goos | | | | Abolnik,C.; Rauff,D.L.; Pieterse,R.; Peyrot,B.; Choma,P.; Ebersohn,K.; Phiri,T. | | | 2017-07-13 | | 2018-04-04 |
| EPI_ISL_387135 | EPI1575305\|A | EPI1575288\|A/d | EPI1575338\|A/duc | EPI1575347\|A/duc | EPI1575302\|A/duc | EPI1575315\|A/duc | EPI1575309\|A/duc | EPI1575283\|A/duc | A/duck/Egypt/N13736E/2017 | A / H5N8 | | Africa / Egypt | Anas sp. | | | | Kandeil,A.; Webby,R.; McKenzie,P.; Kayali,G.; Ali,M.A. | | | 2017-02-15 | | 2018-06-19 |
| EPI_ISL_387135 | EPI1575305\|A | EPI1575288\|A/d | EPI1575338\|A/duc | EPI1575347\|A/duc | EPI1575302\|A/duc | EPI1575315\|A/duc | EPI1575309\|A/duc | EPI1575283\|A/duc | A/duck/Egypt/N13736E/2017 | A / H5N8 | | Africa / Egypt | Anas sp. | | | | Kandeil,A.; Webby,R.; McKenzie,P.; Kayali,G.; Ali,M.A. | | | 2017-02-15 | | 2018-06-19 |
| EPI_ISL_387138 | EPI1575290\|A | EPI1575326\|A/te | EPI1575286\|A/tea | EPI1575319\|A/tea | EPI1575289\|A/tea | EPI1575312\|A/tea | EPI1575318\|A/tea | EPI1575292\|A/tea | A/teal/Egypt /823C/2016 | A / H5N8 | | Africa / Egypt | Anas crecca | | | | Kandeil,A.; Webby,R.; McKenzie,P.; Kayali,G.; Ali,M.A. | | | 2016-12-01 | | 2018-06-19 |
| EPI_ISL_387139 | EPI1575293\|A | EPI1575325\|A/d | EPI1575306\|A/duc | EPI1575344\|A/duc | EPI1575341\|A/duc | EPI1575300\|A/duc | EPI1575332\|A/duc | EPI1575291\|A/duc | A/duck/Egypt/F13666A/2017 | A / H5N8 | | Africa / Egypt | Anas sp. | | | | Kandeil,A.; Webby,R.; McKenzie,P.; Kayali,G.; Ali,M.A. | | | 2017-01-26 | | 2018-06-19 |
| EPI_ISL_387140 | EPI1575336\|A | EPI1575351\|A/te | EPI1575348\|A/tea | EPI1575353\|A/tea | EPI1575345\|A/tea | EPI1575324\|A/tea | EPI1575294\|A/tea | EPI1575320\|A/tea | A/teal/Egypt/1202C/2017 | A / H5N8 | | Africa / Egypt | Anas crecca | | | | Kandeil,A.; Webby,R.; McKenzie,P.; Kayali,G.; Ali,M.A. | | | 2017-02-25 | | 2018-06-19 |
| EPI_ISL_387140 | EPI1575336\|A | EPI1575351\|A/te | EPI1575348\|A/tea | EPI1575353\|A/tea | EPI1575345\|A/tea | EPI1575324\|A/tea | EPI1575294\|A/tea | EPI1575320\|A/tea | A/teal/Egypt/1202C/2017 | A / H5N8 | | Africa / Egypt | Anas crecca | | | | Kandeil,A.; Webby,R.; McKenzie,P.; Kayali,G.; Ali,M.A. | | | 2017-02-25 | | 2018-06-19 |
| EPI_ISL_389757 | | | | EPI1585023\|A/duc | | EPI1585024\|A/duc | | | A/duck/Egypt/N14200B/2017 | A / H5N8 | E1 | Africa / Egypt | Anas sp. | | | | Kandeil,A.; Webby,R.; McKenzie,P.; Kayali,G.; Ali,M.A. | | | 2017-06-09 | | 2018-10-01 |
| EPI_ISL_389773 | | | | EPI1585055\|A/chi | | EPI1585056\|A/chi | | | A/chicken/Egypt/Q13936B/2017 | A / H5N8 | E1 | Africa / Egypt | Gallus gallus | | | | Kandeil,A.; Webby,R.; McKenzie,P.; Kayali,G.; Ali,M.A. | | | 2017-04-12 | | 2018-10-01 |
| EPI_ISL_399644 | | | | EPI1634518\|A/Du | | EPI1789076\|A/Du | | | A/Duck/Egypt/SMG4/2019 | A / H5N8 | | Africa / Egypt | Duck | | | | Tantawy,A.S.; Sultan,H.; Arafa,A.; Hagag,N. | | | 2019-01-23 | | 2019-11-11 |
| EPI_ISL_400027 | EPI1638999\|A | EPI1638796\|A/T | EPI1638795\|A/Tur | EPI1638797\|A/Tur | EPI1638798\|A/Tur | EPI1638799\|A/Tur | EPI1638800\|A/Tur | EPI1638801\|A/Tur | A/Turkey/Egypt/AI20285/2019 | A / H5N8 | | Africa / Egypt | Turkey | Timm Harder (Friedrich-Loeffler-Institut) | Friedrich-Loeffler-Institut | | Hassan,KE;El-Kady,MF;Harder,T. | Poultry Diseases Department ,Faculty of Veterinary Med | | 2019-04-03 | 2019-12-18 | 2019-12-16 |
| EPI_ISL_400043 | EPI1639000\|A | EPI1638783\|A/C | EPI1638784\|A/Chi | EPI1636681\|A/Chi | EPI1638785\|A/Chi | EPI1638786\|A/Chi | EPI1636740\|A/Chi | EPI1636954\|A/Chi | A/Chicken/Egypt/AI20286/2019 | A / H5N8 | | Africa / Egypt | Chicken | Timm Harder (Friedrich-Loeffler-Institut) | Friedrich-Loeffler-Institut | | Hassan,KE;El-Kady,MF;Harder,T. | Poultry Diseases Department ,Faculty of Veterinary Med | | 2019-04-23 | 2019-12-18 | 2019-12-16 |
| EPI_ISL_404993 | EPI1665421\|A | EPI1665422\|A/w | EPI1665423\|A/whi | EPI1665428\|A/whi | EPI1665425\|A/whi | EPI1665426\|A/whi | EPI1665427\|A/whi | EPI1665424\|A/whi | A/white-fronted goose/Germany-BB/AI00018/2020 | A / H5N8 | | Europe / Gern | White-fronted | Jacqueline King (Friedrich-Loeffler-Institut) | Friedrich-Loeffler-Institut | | | Landeslabor Brandenburg-Berlin | | 2020-01-16 | 2020-01-22 | 2020-01-22 |
| EPI_ISL_405391 | EPI1667595\|A | EPI1667596\|A/cl | EPI1667597\|A/chi | EPI1667598\|A/chi | EPI1667599\|A/chi | EPI1667600\|A/chi | EPI1667601\|A/chi | EPI1667602\|A/chi | A/chicken/Czech Republic/1175-1/2020 | A / H5N8 | | Europe / Czec | Gallus gallus c | Alexander Nagy (State Veterinary Institute Prague) | State Veterinary Institute Prague | 1175/20-1 | Nagy,A | State Veterinary Institute Prague | | 2020-01-17 | 2020-01-24 | 2020-01-24 |
| EPI_ISL_417414 | EPI1718618\|A | EPI1718617\|A/b | EPI1718616\|A/bus | EPI1718611\|A/bus | EPI1718614\|A/bus | EPI1718613\|A/bus | EPI1718612\|A/bus | EPI1718615\|A/bus | A/buzzard/Germany-SN/AI00285/2020 | A / H5N8 | | Europe / Gern | Avian | Jacqueline King (Friedrich-Loeffler-Institut) | Friedrich-Loeffler-Institut | 2020AI00285 | | Landesuntersuchungsanstalt für das Gesundheits- und V | | 2020-03-20 | 2020-05-20 | 2020-03-25 |
| EPI_ISL_419220 | EPI1721661\|A | EPI1721662\|A/tu | EPI1721663\|A/turk | EPI1721664\|A/turk | EPI1721665\|A/turk | EPI1721666\|A/turk | EPI1721667\|A/turk | EPI1721668\|A/turk | A/turkey/Hungary/1020_20VIR749-1/2020 | A / H5N8 | | Europe / Hung | Turkey | Ambra Pastori (Istituto Zooprofilattico Sperimentale Delle Venezie) | Istituto Zooprofilattico Sperimentale Delle Venezie | 1020_20VIR7 | Bálint,Ádám;Malik,Péter;Ursu,Krisztina;Bianca,Zecchin;Gianpi | National Food Chain Safety Office Veterinary Diagnosti | 1020 | 2020-01-09 | 2020-04-02 | 2020-04-01 |
| EPI_ISL_503968 | EPI1780066\|A | EPI1780068\|A/cl | EPI1780070\|A/chi | EPI1780072\|A/chi | EPI1780074\|A/chi | EPI1780076\|A/chi | EPI1780078\|A/chi | EPI1780080\|A/chi | A/chicken/Bulgaria/217_20VIR1724-1/2020 | A / H5N8 | | Europe / Bulg | Chicken | | | | Goujgoulova,G.; Slavcheva,I.; Zecchin,B.; Zamperin,G.; Pastori,A.; Schivo,A.; Fusaro,A.; Terregino,C. | | | 2020-03-09 | | 2020-04-02 |
| EPI_ISL_505399 | EPI1787931\|A | EPI1787951\|A/cl | EPI1787944\|A/chi | EPI1787886\|A/chi | EPI1787924\|A/chi | EPI1787958\|A/chi | EPI1787916\|A/chi | EPI1787938\|A/chi | A/chicken/Al-Sharqia/1822FM/2018 | A / H5N8 | | Africa / Egypt | Chicken | | | | Yehia,N.; Elhussieny,M.; Hagag,N.; Arafa,A.; Mohamed,W. | | | 2018-01 | | 2019-06-17 |
| EPI_ISL_505406 | EPI1787925\|A | EPI1787952\|A/cl | | EPI1787893\|A/chi | EPI1787918\|A/chi | EPI1787961\|A/chi | EPI1787912\|A/chi | EPI1787933\|A/chi | A/chicken/AL Suez/184/2018 | A / H5N8 | | Africa / Egypt | Chicken | | | | Yehia,N.; Elhussieny,M.; Hagag,N.; Arafa,A.; Mohamed,W. | | | 2018-01 | | 2019-06-17 |
| EPI_ISL_505411 | EPI1787927\|A | EPI1787947\|A/T | EPI1787942\|A/Tur | EPI1787898\|A/Tur | EPI1787920\|A/Tur | EPI1787955\|A/Tur | EPI1787917\|A/Tur | EPI1787936\|A/Tur | A/Turkey/Beni Sueif/18296F/2018 | A / H5N8 | | Africa / Egypt | Turkey | | | | Yehia,N.; Elhussieny,M.; Hagag,N.; Arafa,A.; Mohamed,W. | | | 2018-02 | | 2019-06-17 |
| EPI_ISL_505413 | EPI1787928\|A | EPI1787948\|A/cl | EPI1787943\|A/chi | EPI1787900\|A/chi | EPI1787923\|A/chi | EPI1787956\|A/chi | EPI1787911\|A/chi | EPI1787937\|A/chi | A/chicken/Albehra/18319F/2018 | A / H5N8 | | Africa / Egypt | Chicken | | | | Yehia,N.; Elhussieny,M.; Hagag,N.; Arafa,A.; Mohamed,W. | | | 2018-04 | | 2019-06-17 |
| EPI_ISL_505425 | EPI1787930\|A | EPI1787950\|A/D | EPI1787941\|A/Du | | EPI1787922\|A/Du | EPI1787957\|A/Du | EPI1787914\|A/Du | EPI1787934\|A/Du | A/Duck/Cairo/1795CA/2017 | A / H5N8 | | Africa / Egypt | Duck | | | | Yehia,N.; EL hussieny,M.; Hagag,N.; Arafa,A.E.; Mohamed,W. | | | 2017-03 | | 2019-06-17 |
| EPI_ISL_529179 | EPI1802158\|A | EPI1802159\|A/d | EPI1802160\|A/duc | EPI1802161\|A/duc | EPI1802162\|A/duc | EPI1802163\|A/duc | EPI1802164\|A/duc | EPI1802165\|A/duc | A/duck/Netherlands/16014829-001005/2016 | A / H5N8 | E2 | Europe / Neth | Duck | Rene Heutink (Wageningen Bioveterinary Research) | Wageningen Bioveterinary Research | | Beerens, Nancy; Harders, Frank; Verschuren-Pritz, Sylvia; Bo | Wageningen Bioveterinary Research | 16014829-00 | 2016-11-25 | 2020-09-11 | 2020-09-10 |

Supplementary File S3: GISAID Acknowledgements
